# GWAS analysis combined with QTL mapping identify *CPT3* and *ABH* as genes underlying dolichol accumulation in Arabidopsis

**DOI:** 10.1101/2020.10.21.348912

**Authors:** Katarzyna Gawarecka, Joanna Siwinska, Jaroslaw Poznanski, Agnieszka Onysk, Przemyslaw Surowiecki, Liliana Surmacz, Ji Hoon Ahn, Arthur Korte, Ewa Swiezewska, Anna Ihnatowicz

**Affiliations:** Institute of Biochemistry and Biophysics Polish Academy of Sciences, Pawinskiego 5a, 02-106 Warszawa, Poland; Intercollegiate Faculty of Biotechnology of University of Gdansk and Medical University of Gdansk, Abrahama 58, 80-307 Gdansk, Poland; Department of Life Sciences, Korea University, Seoul, Korea; Center for Computational and Theoretical Biology, University of Wurzburg, Emil-Fischer Strasse 32, 97074 Wurzburg, Germany

**Keywords:** isoprenoid, polyprenol, dolichol, natural variation, plant-environment interactions, secondary metabolism, QTL mapping, GWAS

## Abstract

Dolichols (Dols), ubiquitous components of living organisms, are indispensable for cell survival. In plants, as well as other eukaryotes, Dols are crucial for posttranslational protein glycosylation, aberration of which leads to fatal metabolic disorders in humans. Until now, the regulatory mechanisms underlying Dol accumulation remain elusive. In this report, we have analyzed the natural variation of the accumulation of Dols and six other isoprenoids between 120 *Arabidopsis thaliana* accessions. Subsequently, by combining QTL and GWAS approaches, we have identified several candidate genes involved in the accumulation of Dols, polyprenols, plastoquinone, and phytosterols. The role of two genes implicated in the accumulation of major Dols in Arabidopsis – the AT2G17570 gene encoding a long searched for *cis*-prenyltransferase (CPT3) and the AT1G52460 gene encoding an alpha-beta hydrolase (ABH) – is experimentally confirmed. These data will help to generate Dol-enriched plants which might serve as a remedy for Dol-deficiency in humans.

## INTRODUCTION

Isoprenoids (also known as terpenes) are a large and diverse group of compounds comprised of more than 40,000 chemical structures (Bohlmann and Keeling, 2008). Linear polymers containing from 5 to more than 100 isoprene units are called polyisoprenoids (Swiezewska and Danikiewicz, 2005). Due to the hydrogenation status of their OH-terminal, (α-) isoprene unit, polyisoprenoids are subdivided into α-unsaturated polyprenols (hereafter named Prens) and α-saturated dolichols (hereafter named Dols) (Figure 1). Prens are common for bacteria, green parts of plants, wood, seeds, and flowers, while Dols are constituents of plant roots as well as animal and fungal cells (Rezanka and Votruba, 2001). In eukaryotic cells, the dominating polyisoprenoid components are accompanied by traces of their counterparts, e.g., Prens are accompanied by Dols in photosynthetic tissues (Skorupinska-Tudek et al., 2003).

**Figure 1.**
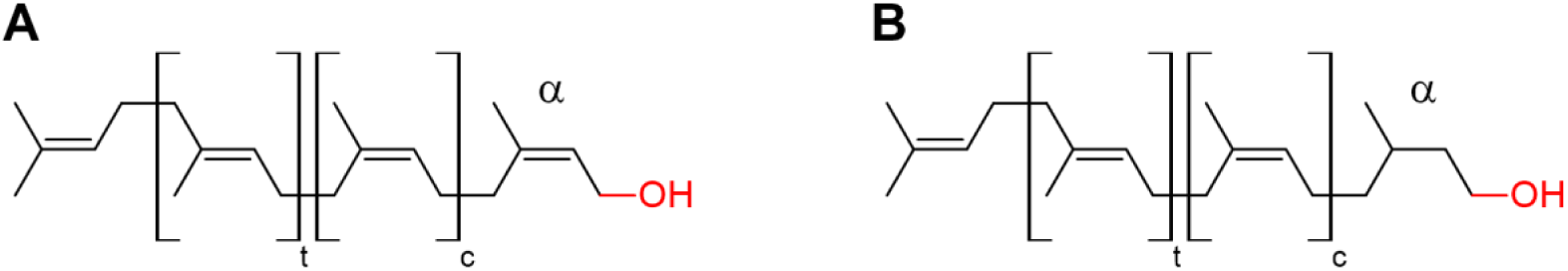
Structures of polyprenol (A) and dolichol (B). t and c stand for the number of internal isoprene units in *trans* and *cis* configuration, respectively. The α-terminal isoprene unit is depicted.

All isoprenoids are synthesized from isopentenyl and dimethylallyl diphosphate (IPP and DMAPP) molecules, which in plants are derived from the cytoplasmic mevalonate (MVA) and plastidial methylerythritol phosphate (MEP) pathways (Hemmerlin et al., 2012; Lipko and Swiezewska, 2016). Formation of the polyisoprenoid chains of both Pren and Dol from IPP is executed by enzymes called *cis*-prenyltransferases (CPTs), which are responsible for elongation of an all-*trans* initiator molecule, most commonly farnesyl or geranylgeranyl diphosphate. This reaction generates a mixture of polyprenyl diphosphates (PolyprenylPP) of similar, CPT-specific, lengths. In *Arabidopsis thaliana* (hereafter named Arabidopsis), only three (Oh et al., 2000; Cunillera et al., 2000; Surowiecki et al., 2019; Kera et al., 2012; Surmacz et al., 2014; Akhtar et al., 2017) out of nine putative CPTs (Surmacz and Swiezewska, 2011) have been characterized at the molecular level. Interestingly, none of these well-characterized CPTs (CPT1, -6 or -7) is responsible for the synthesis of the major ‘family’ of Dols (Dol-16 dominating) accumulated in Arabidopsis tissues.

The polyprenyl diphosphates resulting from CPT activity undergo then either dephosphorylation to Prens and/or reduction to Dols. The reduction reaction is catalyzed by polyprenol reductases, two of which have been recently described in Arabidopsis (Jozwiak et al., 2015). Although most enzymes functioning in the Pren and Dol biosynthesis pathways have been identified, the potential regulatory mechanisms remain unknown.

Isoprenoids are implicated in vital processes in plants, e.g. in photosynthesis and stress response (chlorophylls, carotenoids, plastoquinone, and tocopherols), or in the synthesis of plant hormones (carotenoids, sterols), or they function as structural components of membranes (sterols) (Tholl, 2015). Polyisoprenoids are modulators of the physico-chemical properties of membranes, but they are also involved in other specific processes. Dolichyl phosphate (DolP) serves as an obligate cofactor for protein glycosylation and for the formation of glycosylphosphatidylinositol (GPI) anchors, while Prens, in turn, have been shown to play a role in plant photosynthetic performance (Akhtar et al., 2017). Importantly, an increased content of Prens improves the environmental fitness of plants (Hallahan and Keiper-Hrynko, 2006). Additionally, it has also been suggested that in plants Prens and Dols might participate in cell response to stress since their content is modulated by the availability of nutrients (Jozwiak et al., 2013) and by other environmental factors (xenobiotics, pathogens, and light intensity) (summarized in Surmacz and Swiezewska, 2011). Moreover, the cellular concentration of Prens and Dols is also considerably increased upon senescence (summarized in Swiezewska and Danikiewicz, 2005). These observations suggest that eukaryotes might possess, so far elusive, regulatory mechanisms allowing them to control polyisoprenoid synthesis and/or degradation. Until now, no systematic analysis of the natural variation of polyisoprenoids has been performed for any plant species.

Most traits important in agriculture, medicine, ecology, and evolution, including variation in chemical compound production, are of a quantitative nature and are usually due to multiple segregating loci (Mackay 2001). Arabidopsis is an excellent model for studying natural variation due to its genetic adaptation to different natural habitats and its extensive variation in morphology, metabolism, and growth (Alonso-Blanco et al., 2009; Fusari et al., 2017). Natural variation for many traits has been reported in Arabidopsis, including primary and secondary metabolism (Mitchell-Olds and Pedersen, 1998; Kliebenstein et al., 2001; Sergeeva et al., 2004; Tholl et al., 2005; Keurentjes et al., 2006; Meyer et al., 2007; Lisec et al., 2008; Rowe et al., 2008; Siwinska et al., 2015). Therefore, in this study, we decided to use the model plant Arabidopsis to explore the natural variation of Prens and Dols. Importantly, Arabidopsis provides the largest and best-described body of data on the natural variation of genomic features of any plant species (Kawakatsu et al., 2016; The 1001 Genomes Consortium, 2016). Over 6,000 different Arabidopsis accessions that can acclimate to enormously different environments (Kramer, 2015) have been described so far (Weigel and Mott, 2009).

In order to understand the genetic basis underlying the variation in polyisoprenoid content, and to identify genes that are responsible for it, we used both a quantitative trait loci (QTL) mapping approach and genome-wide association studies (GWAS). So far, neither QTL nor GWAS has been used for the analysis of Prens and Dols. Traditional linkage mapping usually results in detection of several QTLs with a high statistical power, making it a powerful method in the identification of genomic regions that co-segregate with a given trait in mapping populations (Koornneef et al., 2004; Korte and Farlow, 2013). But the whole procedure including the identification of underlying genes is usually time-consuming and laborious. Moreover, the mapped QTL regions can be quite large, making it sometimes impossible to identify the causative genes. Another issue is that the full range of natural variation is not analyzed in QTL studies using bi-parental populations, because they are highly dependent on the genetic diversity of the two parents and may reflect rare alleles. GWAS studies profit from a wide allelic diversity, high resolution and may lead to the identification of more evolutionarily relevant variation (Kooke et al., 2016).Therefore, it is possible to overcome some limitations of QTL analyses by using the GWAS approach, which can be used to narrow down the candidate regions (Korte and Farlow, 2013; Han et al., 2018). But it should be kept in mind that GWAS also has its limitations, such as dependence on the population structure or the potential for false‐positive errors (Zhu et al., 2008; Korte and Farlow, 2013). We have applied here both QTL mapping and GWAS analyses because it has been shown that the combination of these two methods can alleviate their respective limitations (Zhao et al., 2007; Brachi et al., 2010).

The described here application of QTL and GWAS led to identification of several candidate genes underlying the accumulation of polyisoprenoids. Additionally, to get insight into the biosynthetic pathways of Dols and Prens in a broader cellular context, a set of seven isoprenoid compounds was analyzed and subsequently candidate genes that possibly determine the observed phenotypic variation were selected. The most interesting of the identified genes were *cis-prenyltransferase 3* (*CPT3*, AT2G17570, identified through QTL) and *alpha-beta hydrolase* (*ABH*, AT1G52460, identified through GWAS). CPT3, although biochemically not characterized, has been suggested to possess a CPT-like activity (Kwon et al., 2016), whereas alpha-beta hydrolase has not been previously connected with the regulation of polyisoprenoid biosynthesis. In this work, their involvement in Dol biosynthesis/accumulation is experimentally confirmed. Importantly, identification of CPT3 and ABH described in this study fills the gap in the Dol biosynthetic route in Arabidopsis and makes the manipulation of Dol content in plants feasible. Consequently, an option for the generation of plant tissues with increased Dol content as dietary supplements for individuals suffering from Dol-deficiency is emerging.

## RESULTS

### Phenotypic variation in isoprenoid content among Arabidopsis accessions

A set of 116 natural Arabidopsis accessions, originating from various geographical locations, was carefully selected for a detailed analysis of seven isoprenoid compounds (carotenoids, chlorophylls, Dols, phytosterols, plastoquinone, Prens, and tocopherols). Levels of seven selected isoprenoids were quantified in 3-week-old seedlings grown on solid Murashige-Skoog medium. For all analyzed accessions, the same profiles of isoprenoids were observed, however, their content differed remarkably. Thus, for all accessions, one ‘family’ of Prens composed of 9 to 12 isoprene units (Pren-9 to -12, Pren-10 dominating) and one ‘family’ of Dols (Dol-15 to -18, Dol-16 dominating) were detected (Figure 2); however, the content of Prens and Dols revealed remarkable variation between accessions (Figure 3).

**Figure 2 with 3 supplements.**
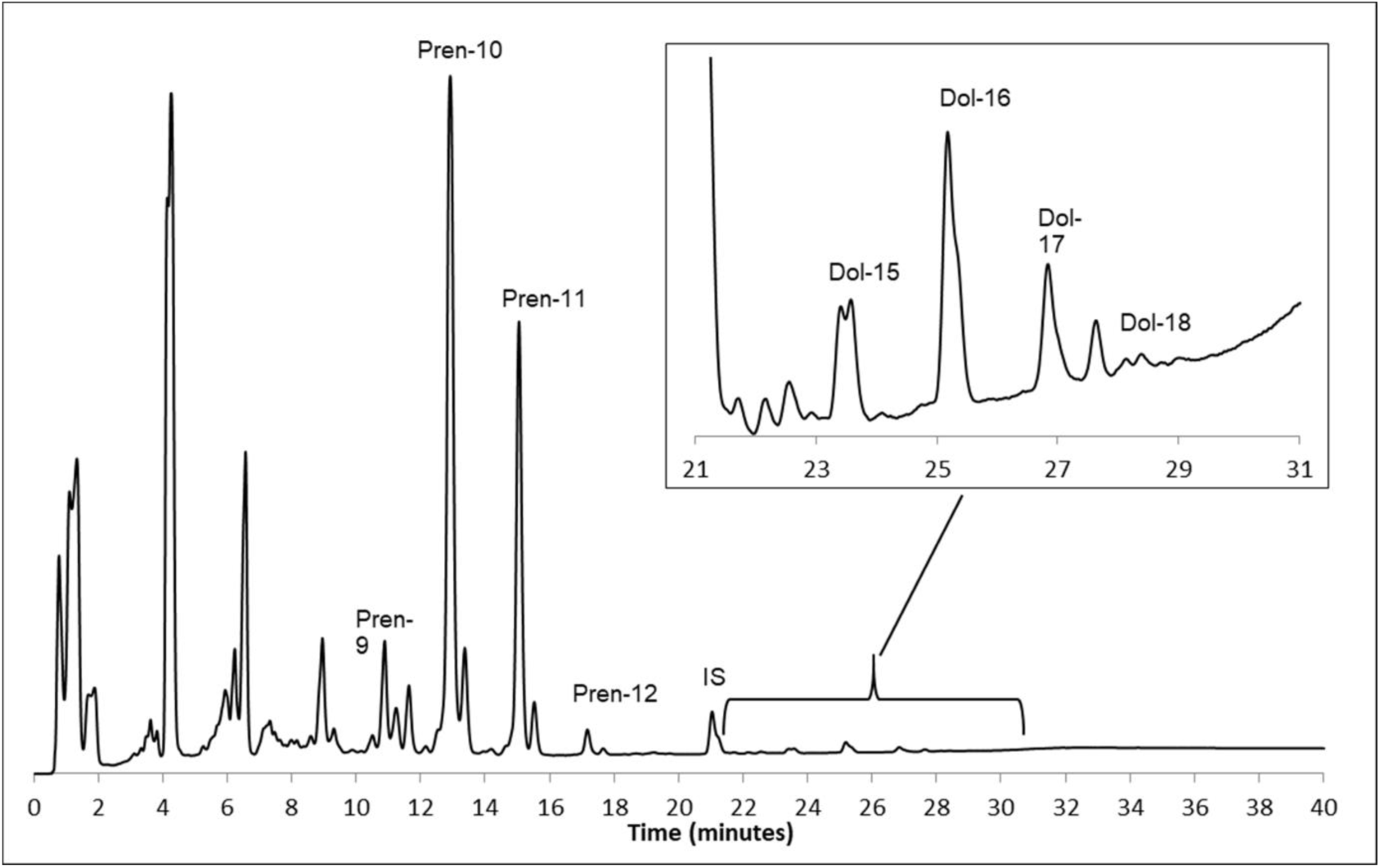
Profiles of polyisoprenoid lipids: polyprenols (Pren) and dolichols (Dol, inset), from Arabidopsis Col-0 seedlings. The same profile of polyisoprenoids was observed for all analyzed accessions. Signals corresponding to Pren-9 to -12 and Dol-15 to -18 were integrated to calculate the total amount of Prens and Dols, respectively. IS indicates the signal of the internal standard (Pren-14) (see Materials and methods and Source Data 1). Profiles of other isoprenoids (phytosterols, plastoquinone and tocopherols) are given in Figure 2-figure supplements 1-3 (see Source Data 2 and 3).

**Figure 3 with 1 supplement.**
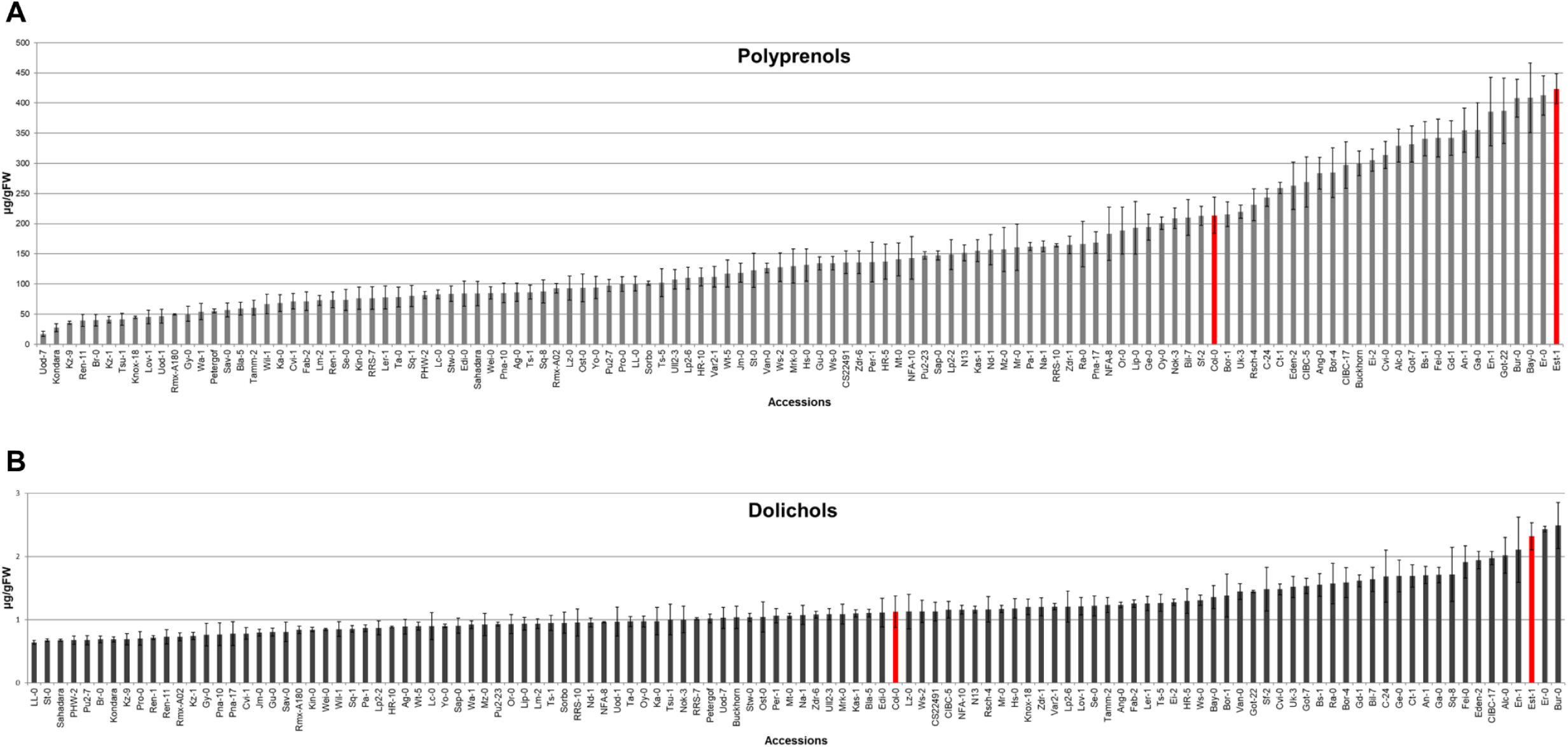
Content of (A) polyprenols (Pren) and (B) dolichols (Dol) in Arabidopsis accessions. Bars representing Col-0 and Est-1 are marked in red. Shown are means ± SD (n=3) (Source Data 4). Content of other isoprenoids (chlorophylls, carotenoids, phytosterols, plastoquinone and tocopherols) in the seedlings of Arabidopsis accessions are given in Figure 3-figure supplement 1 (Source Data 4).

The highest difference in Pren content was observed for the accessions Est-1 and Uod-7 (20-fold), while in Dol content – for LL-0 and Bur-0 (4-fold). Similar observations were noted for the remaining isoprenoids – although the profile was the same for all accessions (Figure 2-figure supplements 1-3), their content revealed substantial differences (Figure 3-figure supplement 1). For phytosterols – 5-fold (Sav-0 vs. Est-1), for plastoquinone – 25-fold (Mr-0 vs. Er-0), for tocopherols – 8-fold (Lip -0 vs. Edi-0), for carotenoids – 4-fold (Est-1 vs. CS22491) and for chlorophylls – 5-fold (Br-0 vs. CS22491) (Figure 3 - figure supplement 1). Detailed analyses revealed considerable differences in the content of 5 out of 7 analyzed compounds (i.e., Prens, Dols, phytosterols, carotenoids, and plastoquinone) between Est-1 and Col-0.

Moreover, Est-1 and Col-0 are the parents of the advanced intercross recombinant inbred lines (AI-RILs) mapping population (EstC), which is an excellent resource for QTL analyses due to a large number of fixed recombination events and the density of polymorphisms (Balasubramanian et al., 2009). For these reasons, the EstC population was selected for further analyses in addition to the analysis of the natural accessions.

### Phenotypic variation in isoprenoid content in the AI-RIL mapping population

Next, the seven isoprenoid compounds described above (carotenoids, chlorophylls, Dols, phytosterols, plastoquinone, Prens, and tocopherols) were quantified in 146 lines of the EstC mapping population and its parental lines (Col-0 and Est-1). The profiles of analyzed isoprenoids were similar to those described above for different accessions, while the content of particular compounds varied among lines of the mapping population (shown in details in Figure 4, Figure 4-figure supplement 1, and Table 1). The range of the content of Prens (Figure 4A), Dols (Figure 4B) and other compounds (Figure 4-figure supplement 1) was broader than that observed for both parental lines, which might suggest that several loci within the EstC population contribute to this phenomenon and it may be explained by the presence of transgressive segregation.

**Table 1.** Isoprenoid content: parental values, ranges, and heritabilities in the AI-RIL mapping population (see Materials and methods and Source Data 5).

| Isoprenoid<br>compound<br>values [$\mu\text{g/gFW}$] | Parental lines | | AI-RILs | | |
| --- | --- | --- | --- | --- | --- |
|  | Col-0 | Est-1 | Range | Median (quartiles) | Heritability<sup>a</sup> |
| Prenols | $116 \pm 10$ | $179 \pm 5$ | 60 – 209 | 129 (104; 153) | 0.55 |
| Dolichols | $0.9 \pm 0.1$ | $1.6 \pm 0.5$ | 0.7 – 2.0 | 1.1 (0.9; 1.2) | 0.55 |
| Chlorophylls | $503 \pm 29$ | $250 \pm 8$ | 222 – 604 | 392 (349; 441) | 0.42 |
| Carotenoids | $125 \pm 18$ | $75 \pm 8$ | 57 – 140 | 94 (84; 104) | 0.43 |
| Phytosterols | $98 \pm 11$ | $125 \pm 3$ | 74 – 154 | 107 (97; 117) | 0.33 |
| Plastoquinone | $99 \pm 12$ | $148 \pm 12$ | 50 – 176 | 111 (97; 127) | 0.47 |
| Tocopherols | $138 \pm 34$ | $226 \pm 36$ | 76 – 288 | 142 (121; 163) | 0.57 |

<sup>a</sup>Measure of total phenotypic variance attributable to genetic differences among genotypes (broad sense heritability) calculated as $V_G/(V_G+V_E)$ .

**Figure 4 with 1 supplement.**
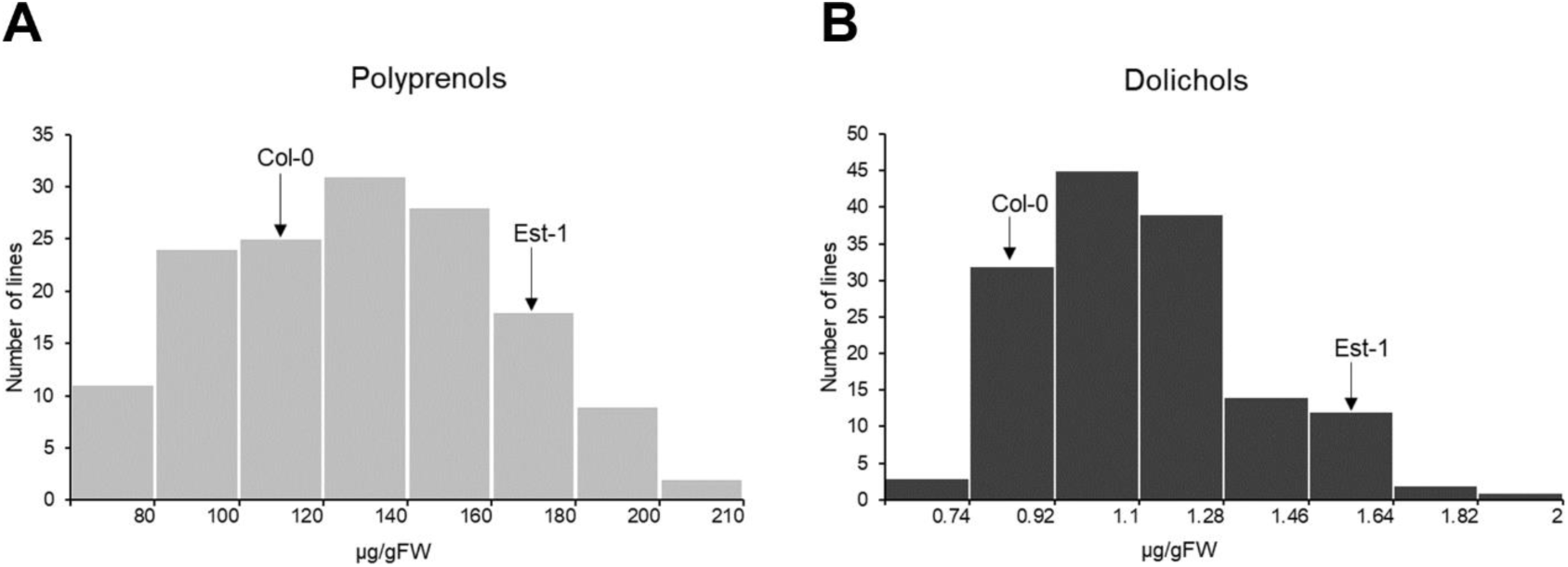
Frequency distribution of the content of polyprenols (A) and dolichols (B) in the seedlings of AI-RILs and their parental lines, Col-0 and Est-1. (see Source Data 5). Each bar covers the indicated range of a particular isoprenoid compound. Frequency distribution of the content of other isoprenoids (chlorophylls, carotenoids, phytosterols, plastoquinone and tocopherols) are given in Figure 4-figure supplement 1 (Source Data 5).

### Estimation of the heritability of isoprenoid levels

To identify the fraction of the observed variation that is genetically determined and whether it can be potentially mapped into QTLs, we estimated the broad sense heritability (*H*^*2*^) for each isoprenoid (Table 1) as described in the Material and Methods section. In the AI-RIL population, the broad sense heritability ranged from 0.33 (for Phytosterols) to 0.55 (for Pren and Dol) and 0.57 (for Tocopherols) (Table 1).

### Identification of QTLs for the accumulation of Dols, Prens, chlorophylls, and carotenoids

The collected biochemical data for the EstC mapping population were subsequently used to map QTL regions underlying the observed phenotypic variation in isoprenoid accumulation. We were able to map QTLs for four types of compounds (Prens, Dols, chlorophylls, and carotenoids). We detected three QTLs on chromosome 5 for Pren accumulation (Figure 5A) (127.3-133.4 cM, 166.5-170.8 cM, and 171.1-173.3 cM), explaining approximately 33% of the phenotypic variance explained (PVE) by these QTLs containing 948 loci (Table 2). For Dol, we detected a QTL region on chromosome 2 (Figure 5B) (64.8-74.4 cM) containing 308 loci (Table 2), which explains approximately 16.8% of the PVE.

**Table 2.**
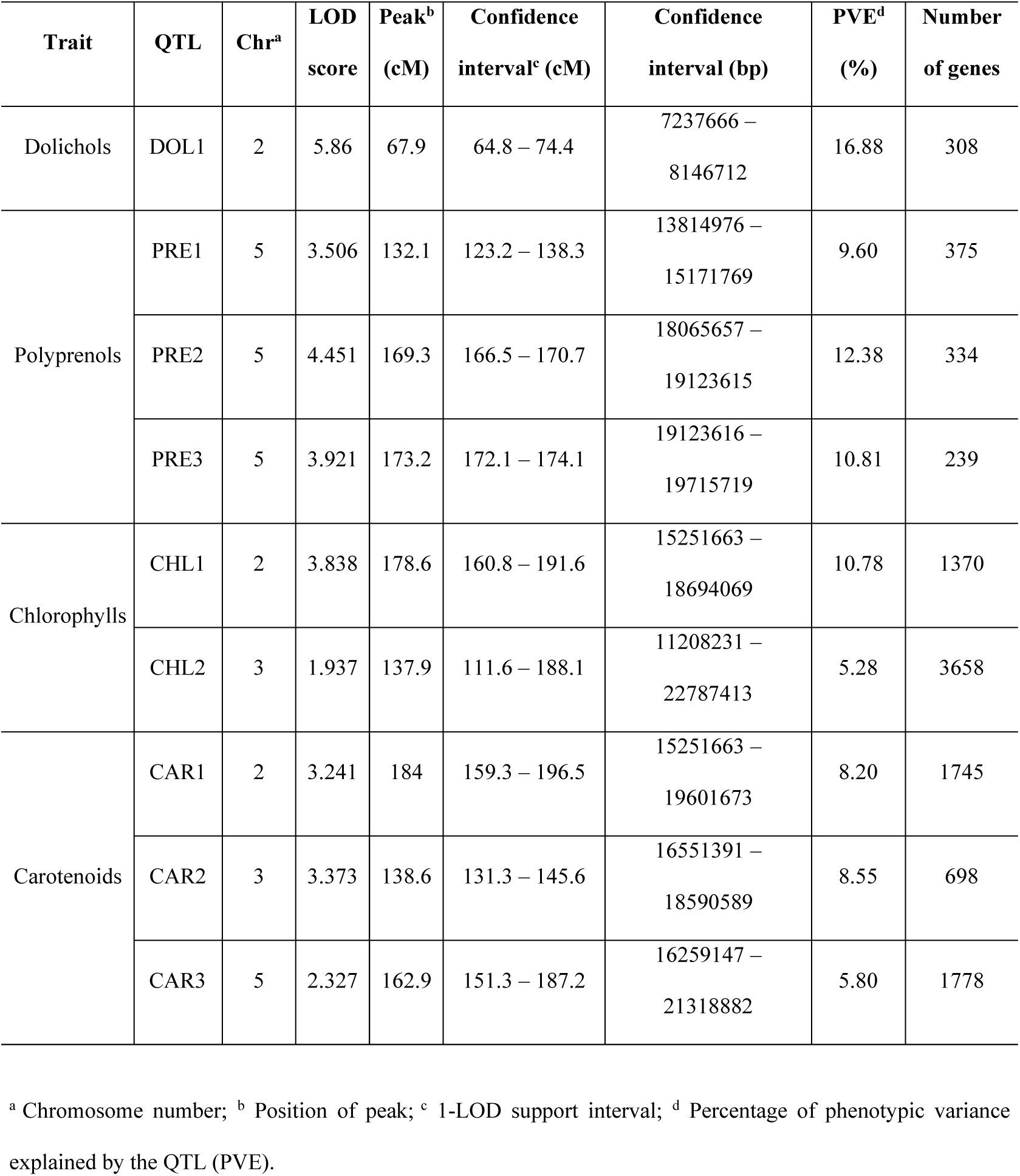
Characteristics of the detected QTLs underlying polyprenol (Pren), dolichol (Dol), chlorophyll and carotenoid accumulation in the AI-RIL population (see Materials and methods and Source Data 6-9).

| Trait | QTL | Chr<sup>a</sup> | LOD score | Peak<sup>b</sup> (cM) | Confidence interval<sup>c</sup> (cM) | Confidence interval (bp) | PVE<sup>d</sup> (%) | Number of genes |
| --- | --- | --- | --- | --- | --- | --- | --- | --- |
| Dolichols | DOL1 | 2 | 5.86 | 67.9 | 64.8 – 74.4 | 7237666 – 8146712 | 16.88 | 308 |
| Polyprenols | PRE1 | 5 | 3.506 | 132.1 | 123.2 – 138.3 | 13814976 – 15171769 | 9.60 | 375 |
|  | PRE2 | 5 | 4.451 | 169.3 | 166.5 – 170.7 | 18065657 – 19123615 | 12.38 | 334 |
|  | PRE3 | 5 | 3.921 | 173.2 | 172.1 – 174.1 | 19123616 – 19715719 | 10.81 | 239 |
| Chlorophylls | CHL1 | 2 | 3.838 | 178.6 | 160.8 – 191.6 | 15251663 – 18694069 | 10.78 | 1370 |
|  | CHL2 | 3 | 1.937 | 137.9 | 111.6 – 188.1 | 11208231 – 22787413 | 5.28 | 3658 |
| Carotenoids | CAR1 | 2 | 3.241 | 184 | 159.3 – 196.5 | 15251663 – 19601673 | 8.20 | 1745 |
|  | CAR2 | 3 | 3.373 | 138.6 | 131.3 – 145.6 | 16551391 – 18590589 | 8.55 | 698 |
|  | CAR3 | 5 | 2.327 | 162.9 | 151.3 – 187.2 | 16259147 – 21318882 | 5.80 | 1778 |

<sup>a</sup> Chromosome number; <sup>b</sup> Position of peak; <sup>c</sup> 1-LOD support interval; <sup>d</sup> Percentage of phenotypic variance explained by the QTL (PVE).

**Figure 5.**
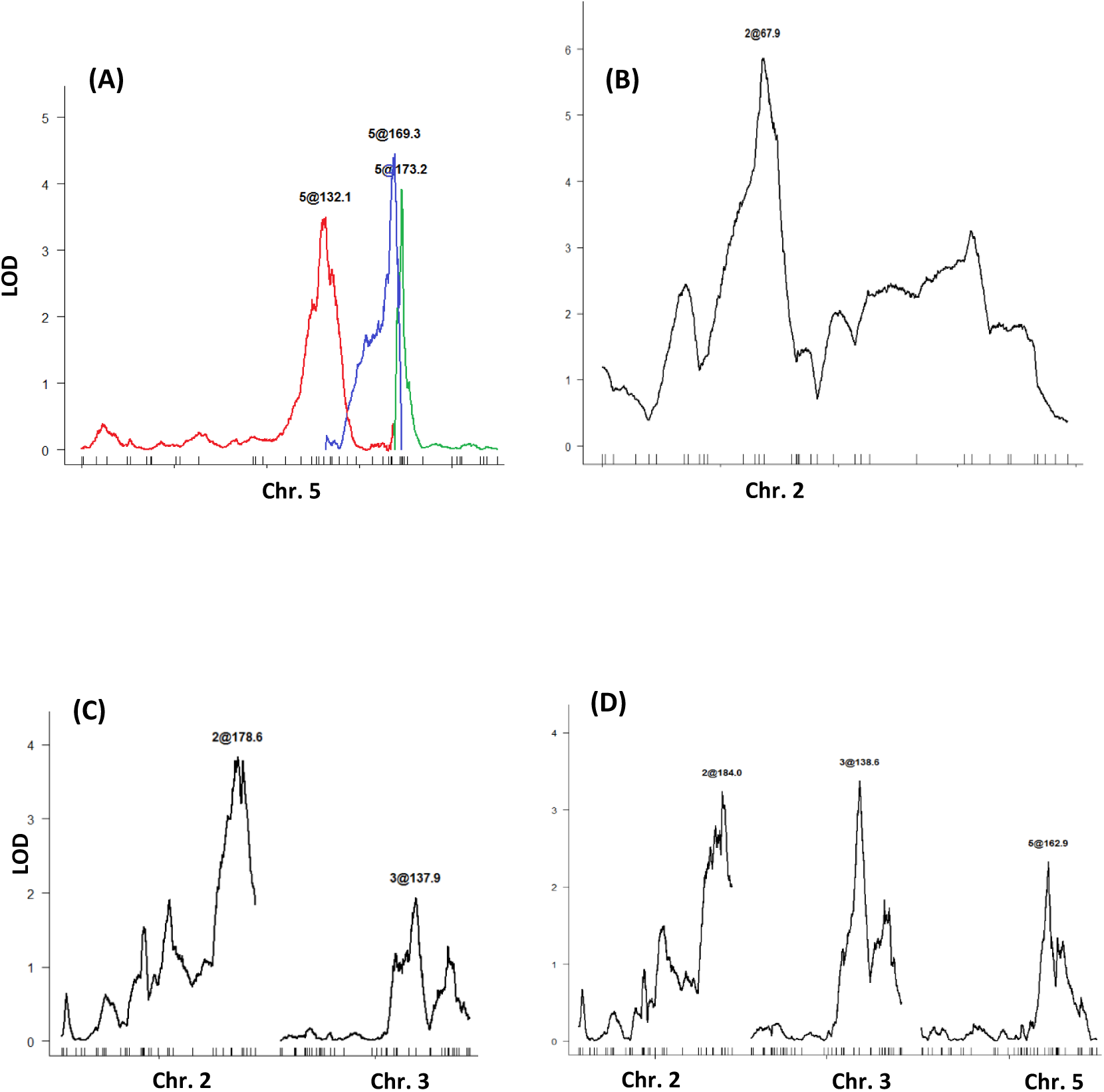
LOD profiles for QTLs underlying the accumulation of selected isoprenoids in the AI-RILs: (A) polyprenols, (B) dolichols, (C) chlorophylls, and (D) carotenoids (see Materials and methods and Source Data 6-9).

Two QTLs were detected for chlorophyll accumulation on chromosome 2 (160.8-191.6 cM) and 3 (111.6-188.1 cM) (Figure 5C), which together explain 16% of the PVE (Table 2). On chromosome 2, 3, and 5 (159.3-196.5 cM, 131.3-145.6 cM, and 151.3-187.2 cM, respectively) (Figure 5D) we identified three QTLs underlying the variation in carotenoid accumulation, as the whole model explains together almost 24% of the PVE (Table 2). It should be underlined that the QTL on chromosome 3 (for chlorophylls) and the QTL on chromosome 5 (for carotenoids) were included in this analysis despite the fact that their LOD scores were slightly below the threshold (below 3) (Figure 5C and Figure 5D, respectively). Interestingly, two of the QTLs identified for chlorophylls and carotenoids, localized on chromosomes 2 and 3, were overlapping.

Our search also revealed two small QTL regions for phytosterols (data not shown); however, they were not analyzed further due to their statistical insignificance (LOD < 3.0). Despite the large set of numerical data, no QTLs were identified for plastoquinone or tocopherols. This might indicate that the mapping population used in this study was not appropriate for investigating these metabolites.

### Selection of candidate genes from QTL mapping

In order to select and prioritize positional candidate genes from the QTL confidence intervals, we conducted a literature screen and an *in silico* analysis (explained in more detail in the Materials and Methods section) that were based on functional annotations, gene expression data and tissue distribution of the selected genes. We analyzed loci from the Dol-associated QTL (DOL1) and from the three Pren-associated QTLs (PRE1, PRE2, PRE3). We selected the intervals that were characterized by the highest percentage of phenotypic variance related to each QTL and the highest LOD score values linked with the lowest number of loci (Table 2). As a result of the above-described procedure of selection and prioritization, we generated four sets of genes – three for Prens (Supplementary file 1) and one for Dol (Supplementary file 2).

Within a set of potential candidate genes for Pren regulation (Supplementary file 1), there was the AT5G45940 gene encoding the Nudix hydrolase 11 (Kupke et al., 2009) with putative IPP isomerase activity. For the regulation of Dol biosynthesis, we identified three loci that might be directly implicated in the process: AT2G17570, encoding a *cis*-prenyltransferase 3 (CPT3), AT2G17370, encoding HMGR2 (hydroxymethylglutaryl Coenzyme-A reductase 2, also called HMG2, a key regulator of the MVA pathway), and AT2G18620, encoding a putative GGPPS2 (geranylgeranyl diphosphate synthase 2). A brief comment on the putative role of the two latter genes in the Dol pathway is presented in Supplementary file 2, while an in-depth characteristics of AT2G17570 (CPT3) is presented below.

### The role of CPT3 in Dol synthesis in Arabidopsis – genetic and biochemical studies

Remarkably, the CPT responsible for the formation of the hydrocarbon backbone of the major Dols (Dol-15 to Dol-17) accumulated in Arabidopsis has not been identified yet. The AT2G17570 gene encoding CPT3 (sometimes named CPT1 (Kera et al., 2012)) is ubiquitously expressed in Arabidopsis organs and, among all nine AtCPTs, it is by sequence homology the closest counterpart of the yeast CPTs that synthesize Dols (Surmacz and Swiezewska, 2011). Preliminary *in vitro* studies revealed that *CPT3*, when co-expressed with *LEW1*, was capable of rescuing the growth defect of a yeast strain devoid of both yeast CPTs: *rer2Δ srt1Δ*, and a thus obtained yeast transformant was able to incorporate a radioactive precursor into polyisoprenoids, although their profile had not been presented (Kwon et al., 2016).

No T-DNA insertion mutant in the *CPT3* gene is available from the NASC collection. For this reason, to analyze *in planta* the involvement of CPT3 in Dol formation, four independent RNAi lines targeting CPT3 for mRNA knockdown (RNAi-1, -12, -14 and -23) and a transgenic line overexpressing CPT3 (OE-7) were generated. The expression level of *CPT3* and the polyisoprenoid content were examined in 4-week-old leaves of these mutants. qRT-PCR analyses revealed that the *CPT3* transcript is significantly reduced (by 40-50%) in the four RNAi lines, and it is nearly 5-fold elevated in the OE line, in comparison to wild-type plants (Figure 6A).

**Figure 6.**
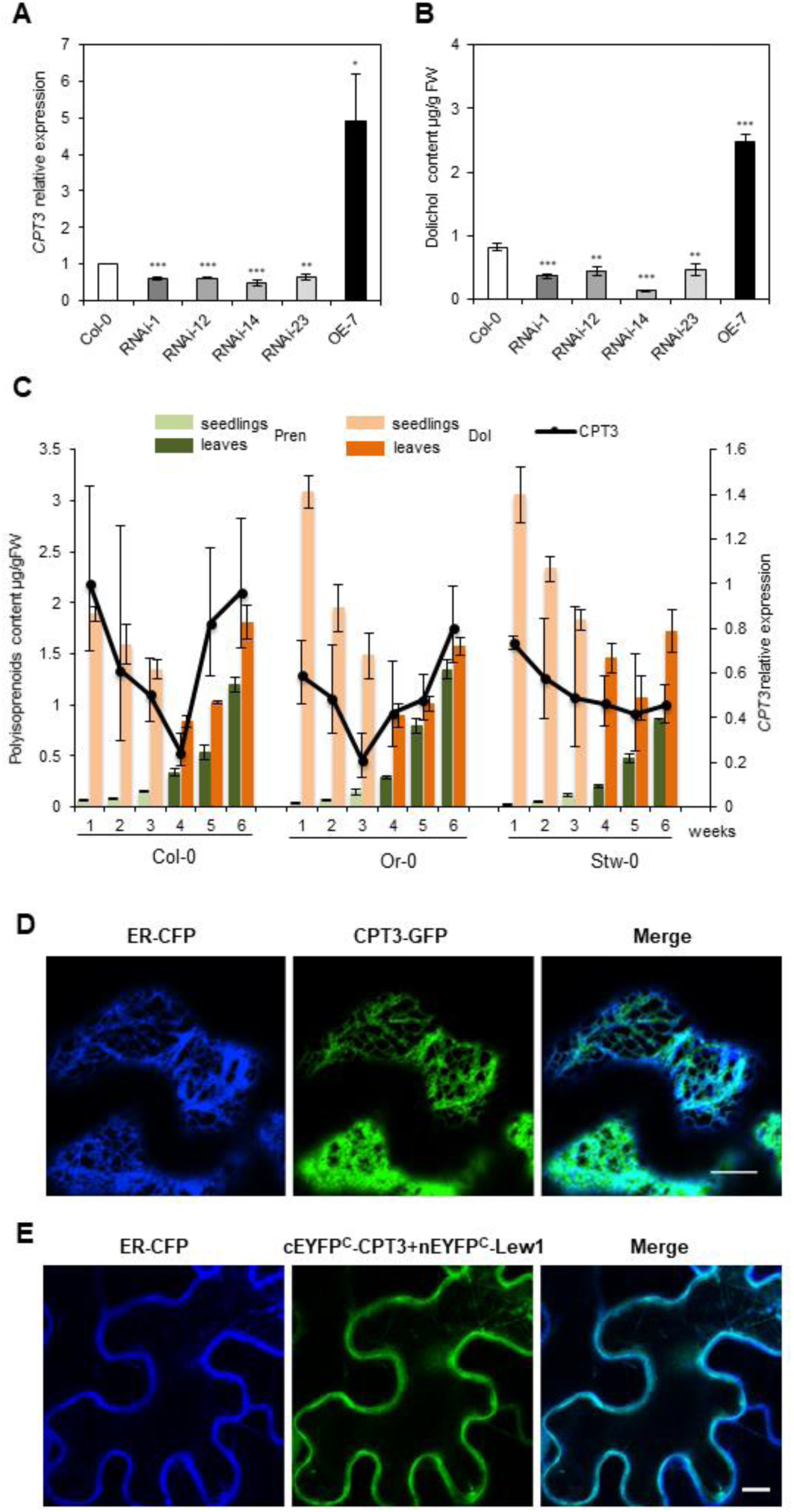
Role of CPT3 in Dol biosynthesis – studies *in planta*. **(A)** Relative expression of *CPT3* and **(B)** content of dolichols (Dol-15 to Dol-17) in the leaves of 4-week-old Arabidopsis plants, measured for wild-type Col-0, four independent RNAi lines targeting *CPT3* (RNAi-1, -12, -14 and -23) and a *CPT3*-overexpressing line (OE-7). The results are means (±SD) of three independent experiments. Asterisks indicate statistically significant differences between WT and mutant plants (* 0.01<P<0.05, ** 0.001<P<0.01, *** P<0.001, Student’s t-test) **(C)** Changes in the levels of *CPT3* mRNA (black curves) and in the content of Dols and Prens (respectively, orange and green bars) in the tissues of three Arabidopsis accessions: Col-0, Or-0, and Stw-0, during the plant life-span. Transcript levels and lipid content were estimated in Arabidopsis seedlings (1-3 weeks, bright colors) and leaves (4-6 weeks, deep colors), shown are means ± SD (n=3). Please note that the content of Prens is rescaled (0.01 multiples are presented) due to their high cellular level. **(D)** Co-localization of fluorescence signals of CPT3-GFP (green) and ER-CFP (compartmental marker, blue) upon transient expression in *Nicotiana benthamiana* leaves. Bar: 10 μm. **(E)** Analysis of CPT3 and Lew1 protein–protein interaction using split yellow fluorescent protein (YFP) BiFC assay in tobacco leaf cells. Shown is the co-localization of fluorescence signals from the CPT1/Lew1 complex (green) and from the compartmental marker ER-CFP (blue). Bar: 10 μm. See Materials and methods and Source Data 10-17.

No visible phenotypic changes were observed between wild type plants and the studied mutant lines under standard growth conditions (data not shown). In contrast, HPLC/UV analysis of total polyisoprenoids revealed a significant decrease in dolichol (Dol-15 – Dol-17, dominating Dol-16) accumulation in *CPT3* RNAi lines – to approx. 50% of the WT for three lines (RNAi-1, -12, and -23) and to approx. 80% for RNAi-14. Not surprisingly, *CPT3-OE* plants accumulated significantly higher amounts of dolichols, reaching 300% of the WT levels (Figure 6B). These results clearly suggest that CPT3 is involved in the biosynthesis of the major family of Dols in Arabidopsis. In line with this, we observed a positive correlation between the level of *CPT3* transcript and the content of Dol during plant development for three of the selected accessions (Figure 6C). This further supports the role of CPT3 in Dol formation; interestingly, no such correlation was noted for Prens (Figure 6C).

CPT3, similarly to numerous other eukaryotic CPTs engaged in Dol biosynthesis (Grabińska et al., 2016), is located in the endoplasmic reticulum (ER), as documented by confocal laser microscopy – in transiently transformed *N. benthamiana* leaves the fluorescence signal of CPT3-GFP fully overlapped with that of the ER marker ER-CFP (Figure 6D).

Finally, functional complementation of the yeast mutant *rer2*Δ by Arabidopsis *CPT3* followed by an analysis of the polyisoprenoid profile of transformants (Figure 7) revealed that solely co-expression of *CPT3* and *LEW1* (Arabidopsis homologue of mammalian NgBR and yeast Nus1, an accessory protein required for activity of some eukaryotic CPTs, Grabińska et al., 2016) resulted in the synthesis of the major family of Dols (Dol-14 to Dol-16, Dol-15 dominating, Figure 7A). Moreover, in line with the cellular function of Dol as an obligate cofactor of protein *N*-glycosylation, only simultaneous expression of *CPT3* and *Lew1* fully rescued the defective glycosylation of the marker protein CPY in *rer2*Δ mutant cells (Figure 7B).

**Figure 7.**
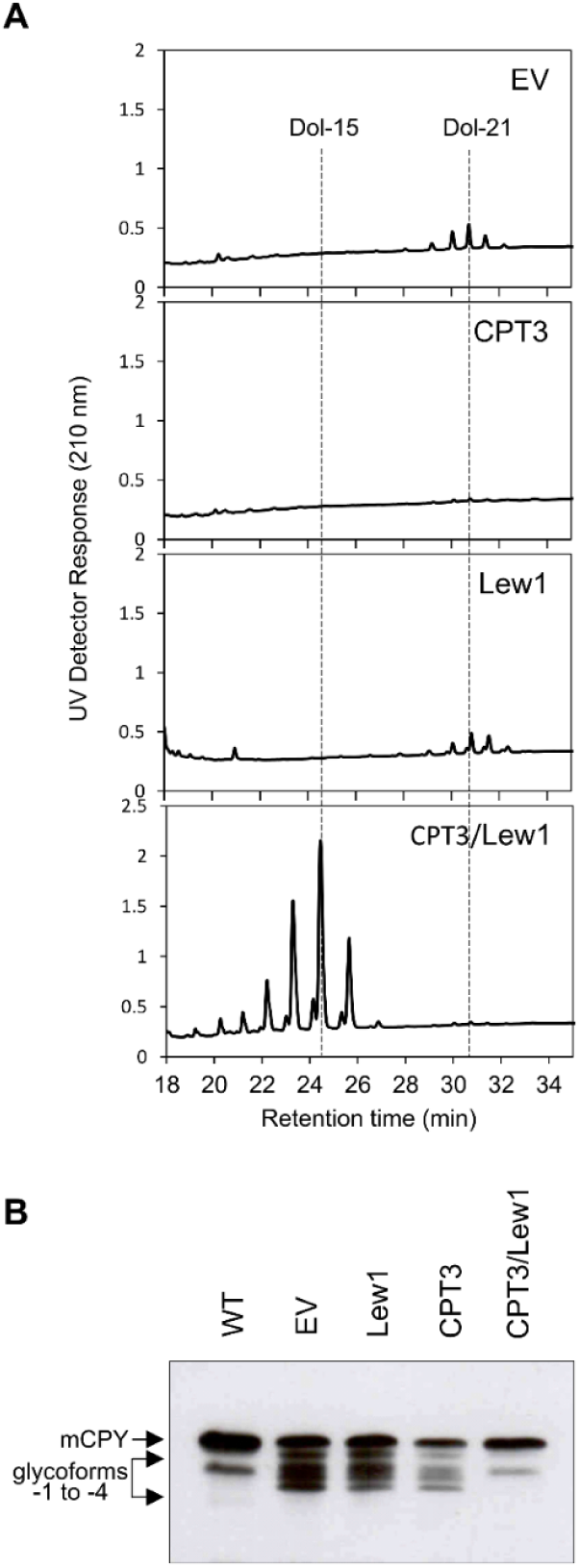
Role of CPT3 in Dol biosynthesis – functional assay in yeast. *AtCPT3* was expressed in a *Saccharomyces cerevisiae* mutant strain devoid of Rer2 activity. **(A)** Polyisoprenoid profiles and **(B)** glycosylation status of CPY analyzed for *rer2*Δ transformed with empty vector, *CPT3, LEW1*, and *CPT3/LEW1*. Representative HPLC/UV chromatograms are shown. The positions of mature CPY (mCPY) and its hypoglycosylated glycoforms (lacking between one and four *N*-linked glycans, -1 to -4) are indicated. See Materials and methods and Source data 18-19.

The physical interaction of CPT3 with Lew1 was confirmed *in planta* using a bimolecular fluorescence complementation (BiFC) assay. nEYFP-C1/CPT3 was transiently co-expressed with cEYFP-N1/Lew1 in *N. benthamiana* leaves. The signal of the enhanced yellow fluorescent protein (EYFP) was localized in the ER (Figure 6E).

Taken together, the genetic and biochemical data presented here clearly show that Arabidopsis CPT3 is a functional ortholog of yRer2 and this verifies the QTL mapping by demonstrating that CPT3 is responsible for Dol synthesis in Arabidopsis.

### Genetic analyses of the variations in metabolite levels in natural accessions - GWAS

As a following step, we used a multi-trait mixed model (Korte et al., 2012) to calculate the genetic correlations between the different traits studied (see Supplementary file 3). Here, we found a strong correlation for the four traits – Prens, phytosterols, plastoquinone, and Dols, which argues for a common genetic correlation of these four traits, and at the same time it shows that they have a negative genetic correlation with the remaining three traits, namely tocopherols, chlorophylls, and carotenoids.

Next, we used the mean phenotypic values of the 116 natural Arabidopsis accessions per trait to perform GWAS. Eighty-six of these lines have been recently sequenced as part of the 1,001 genomes project and full sequence information is readily available (1001 Genomes Consortium, 2016). For the remaining accessions, high-density SNP data have been published earlier (Horton et al., 2012). We used an imputed SNP dataset that combined both sets and has been published earlier (Togninalli et al., 2008). This data set contains ∼ 4 million polymorphisms that segregate in the analyzed accessions. Two million polymorphisms, which had a minor allele count of at least 5, were included in the analysis. At a 5% Bonferroni corrected significance threshold of 2.4 ^*^10^^^-8, significant associations have been found only for three of the seven different compounds analyzed (Dols, plastoquinone, and phytosterols), while no significant associations have been found for the other four compounds (chlorophylls, carotenoids, Prens, and tocopherols). In summary, 2, 7, and 5 distinct genetic regions were significantly associated with Dols, plastoquinone, and phytosterols, respectively. One region on chromosome 1 is found for all three traits. The respective Manhattan plots are shown in Figure 8 and Figure 8-figure supplement 1.

**Figure 8 with 1 supplement.**
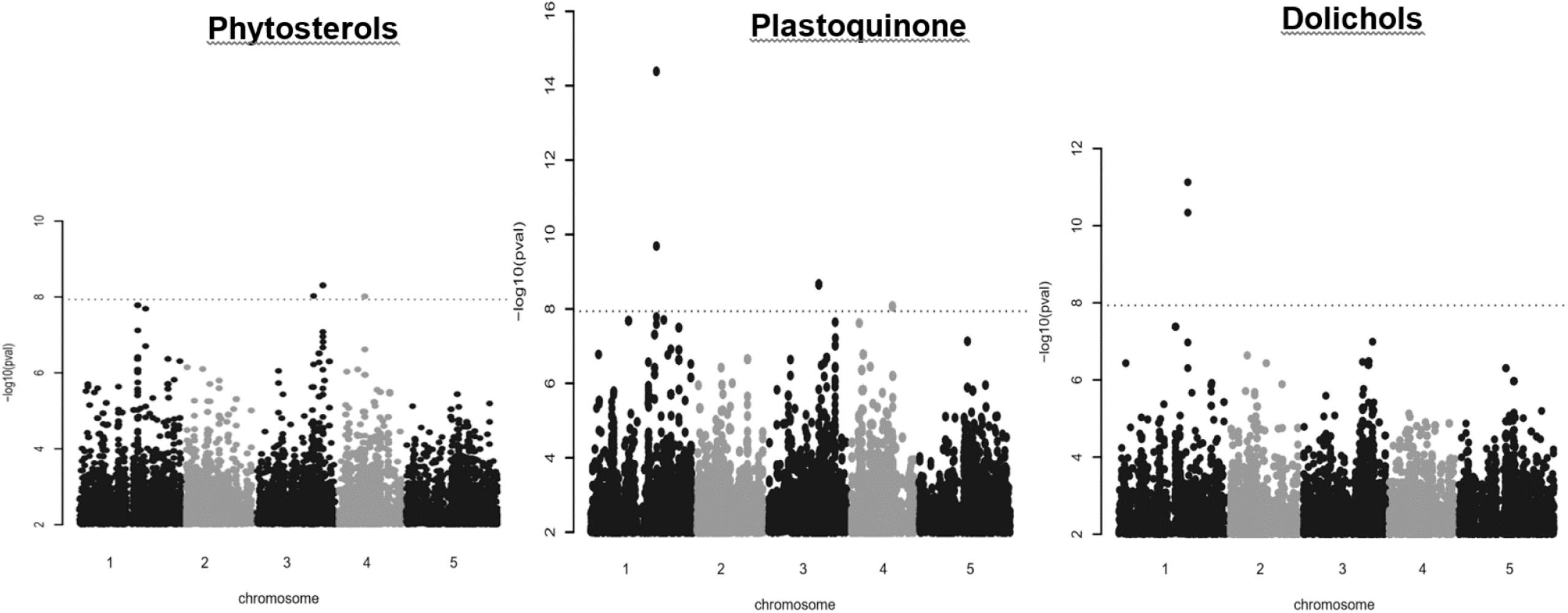
Manhattan plot of genome-wide associations for phytosterols, plastoquinone, and dolichols. The dotted horizontal lines indicate a significance level of 0.05 after Bonferroni correction for multiple testing (see Materials and methods and Source Data 20). Manhattan plot of genome-wide association results for polyprenols, chlorophylls and tocopherols are shown in Figure 8-figure supplement 1 (Source Data 20).

Summarizing, we found 4 SNPs, representing two different regions, that were associated with Dol content. The first of the associated polymorphisms is at position 19,545,459 on chromosome 1 and it codes for a non-synonymous AA-exchange (Q270K) in the first exon of AT1G52450, a gene involved in ubiquitin-dependent catabolic processes. The second polymorphism is located at position 19,540,865: it is upstream of AT1G52450 and in the 3′ UTR of the neighboring gene AT1G52440, which encodes a putative alpha-beta hydrolase (ABH). A second putative ABH (AT1G52460) is also within 10 kb of these associations. The remaining two significant associations are on chromosome 3 (positions 18,558,714 and 18,558,716, respectively) and they code for one non-synonymous (V113G) and one synonymous substitution in an exon of the gene AT3G50050, which encodes the auxin-related transcription factor Myb77 (Shin et al., 2007).

For plastoquinone, 26 SNPs, spread across 7 distinct genomic regions, have been found. The most significant SNP is located at chromosome 1 at position 19,545,459 and is identical to the one reported for Dol. Despite the fact that other significant SNPs are spread over 3 different chromosomes, they are all in linkage disequilibrium (LD) with each other, indicating only one independent association. And indeed, if the lead SNP is added as a cofactor to the model (Segura et al., 2012), none of the remaining SNPs stays above the threshold, indicating only one causative association. It is noteworthy that many of these other SNPs are directly located in transposable elements.

For phytosterols, 10 SNPs at 5 distinct genetic regions showed significant associations. One of these is again the same SNP that has been reported above for plastoquinone and Dols. Interestingly, this polymorphism does not show the strongest association with phytosterols, but three other sequence variants, located around 19.67 Mb on chromosome 3, are in perfect LD and show a stronger association. These polymorphisms are located between AT3G53040, encoding a LEA protein, and AT3G53050, which encodes an enzyme involved in hydrolyzing *O*-glycosyl compounds.

The identification of AT1G52450 and two neighboring genes as putative regulators of the accumulation of Dols, plastoquinone, and phytosterols prompted us to analyze the phenotypes of the respective Arabidopsis T-DNA insertion mutants. Interestingly, a significant increase in the content of Dols (3- and 10-fold, respectively, comparing to control WT plants) was noted for two analyzed heterozygotic AT1G52460-deficient lines: SALK_066806 and GK_823G12. Moreover, in the SALK_066806 line phytosterol content was also increased (167.8 ± 20.3 vs. 117.4 ± 23.2 μg/g of fresh weight) and plastoquinone content was considerably decreased (27.3 ± 2.0 vs 56.7 ± 5.2 μg/g of fresh weight). It is worth noting that mutations in the AT1G52460 gene did not affect the content of Prens – this gene has not come up as a putative Pren regulator (Figure 9). Moreover, these mutant plants developed deformed, curled leaves (Figure 9-figure supplement 1). Other analyzed homozygotic mutants (carrying insertions in the genes AT1G52440 and AT1G52450) did not show significant differences neither in isoprenoid content nor in macroscopical appearance (data not shown).

**Figure 9 with 1 supplement.**
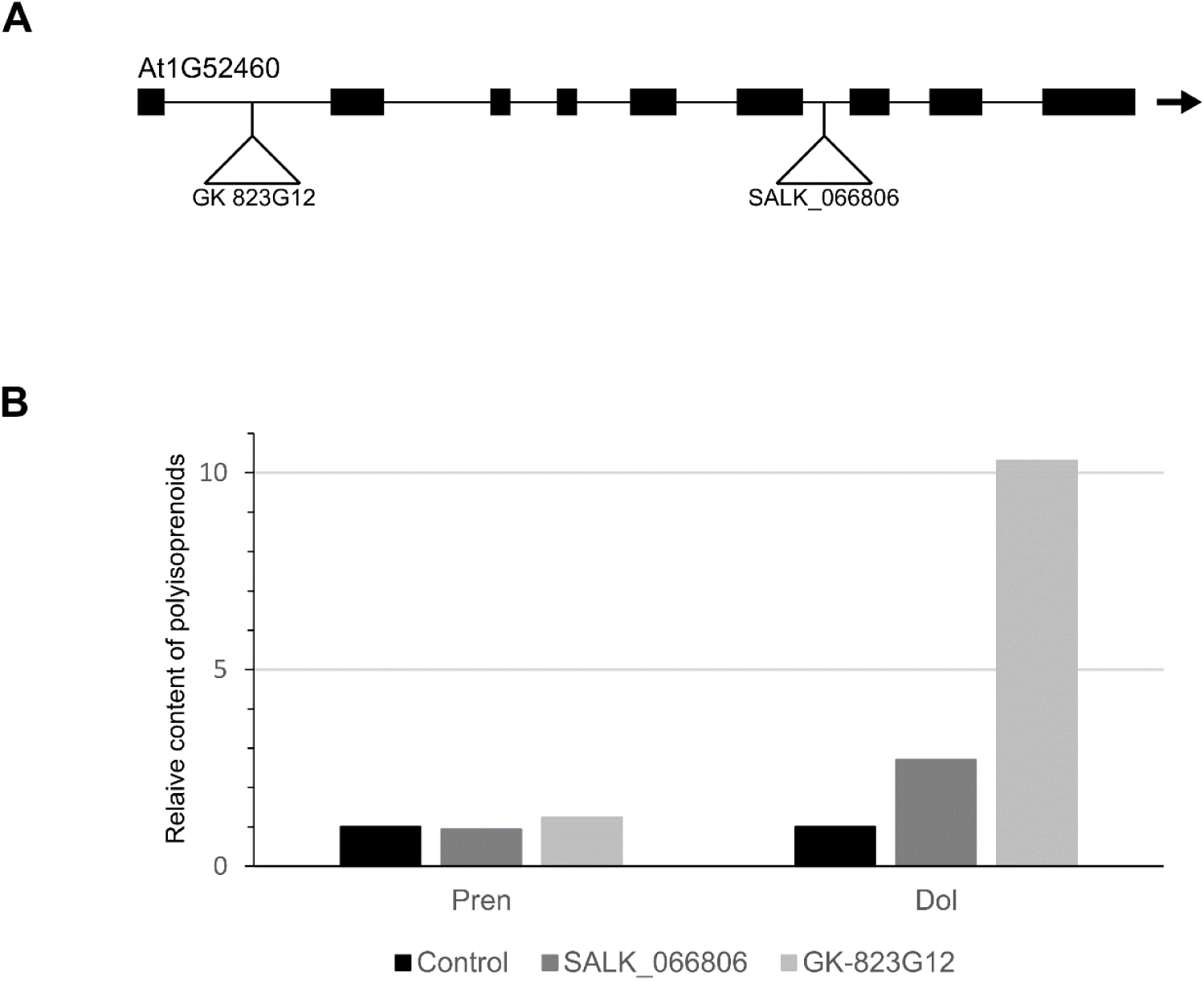
Effect of AT1G52460 deficiency on the content of polyisoprenoids. **(A)** AT1G52460 gene structure, exons and introns are indicated by thick and thin lines, respectively. The T-DNA insertion sites in two independent mutant lines: SALK_066806 and GK_823G12 are depicted. **(B)** The relative content of Dols and Prens estimated in leaves of 3-week-old plants using HPLC/UV (representative results are shown, see Source Data 21). The phenotypic appearance of 4-week-old detached leaves of AT1G52460-deficient line (SALK_066806, *abh* heterozygous mutant) and wild-type (Col-0) plants grown in soil is shown in Figure 9-figure supplement 1.

### Correlation analyses of isoprenoid accumulation in the various accessions and in the mapping population-a statistical meta-analysis

As a final step, we conducted a detailed statistical meta-analysis of the studied traits in the different Arabidopsis accessions and in the lines of the EstC mapping population. Numerous correlations were found for the content of seven isoprenoid compounds estimated in the seedlings of natural accessions and the mapping population (Figure 10A and 10B, respectively). Moreover, we clearly identified some outliers (Grubbs test at significance level α=0.001) (Grubbs 1950). For plastoquinone, seven values corresponding to three accessions (Er-0, Est-1, and Fei-0) were unequivocally assigned as outliers, for carotenoids – three values corresponding to a single accession (Ren-1), for phytosterols a single outlier was identified in the natural accessions and for Dols in the mapping population (Figure10-figure supplement 1). All these outliers, denoted by red triangles in Figure 10, were filtered out in the statistical analysis of metabolite distribution and the correlation analyses (Figure 10A and 10B). For both datasets, the analysis of metabolite correlations revealed the highest correlation for chlorophylls vs. carotenoids (R>0.97), while four other metabolites – phytosterols, Prens, plastoquinone, and Dols – also correlated with each other significantly (p<0.0001). Tocopherol accumulation correlated only occasionally with the other metabolites (Supplementary file 4). Based on the structural similarity between Prens and Dols, some level of similarity between the mechanisms regulating their accumulation might be expected. However, the obtained values for the correlation between Prens and Dols among the tested accessions (0.325, p=0.0001) and among the AI-RILs (0.608, p=0.0001) suggest that some differences exist between these two subgroups of polyisoprenoids.

**Figure 10 with 1 supplement.**
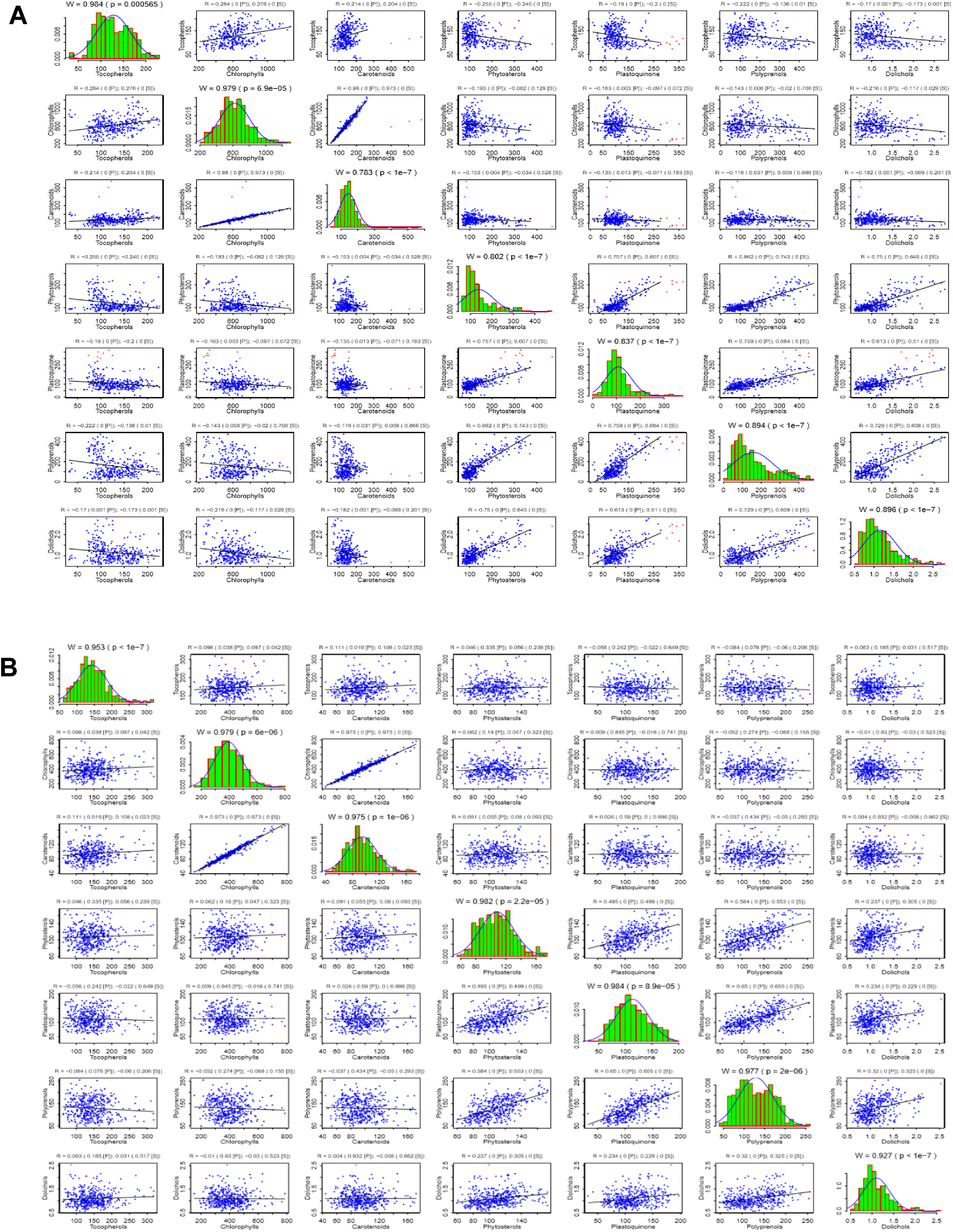
Correlations between the content of seven metabolites estimated in the seedlings of Arabidopsis accessions (A) and the EstC mapping population (B). The original distributions (green bars), together with the approximation of the normal distribution of the data (blue curve) with outliers removed, are presented on the diagonal. Correlation patterns for each metabolite pair are presented at the appropriate intersection; please note that outliers (red dots) were not taken into consideration for the analysis. Above each diagonal panel, the Shapiro-Wilk statistics (W, p) for normal distribution is presented, while for out-of-diagonal panels Pearson (P) and Spearman (S) correlation coefficients together with the associated significance levels are shown (please note that ‘0’ means p< 1e-7). Bearing in mind the statistically significant deviations from normal distribution shown in the diagonal panels, the significance of the observed correlations should be interpreted in terms of the Spearman rather than Pearson coefficient (see Materials and methods and Source Data 22). Cumulative distribution functions (CDF) of the content of seven studied metabolites analyzed in the seedlings of Arabidopsis accessions and AI-RILs are shown in Figure 10-figure supplement 1 (Source Data 22).

Importantly, all the strongest genetic correlations detected for particular metabolites (Supplementary file 3) were also identified as the most significant (p <0.0001) for metabolic data-based analysis and this is valid both for the natural accessions and for the EstC mapping population lines (Supplementary file 4). Moreover, a consistent trend of correlations (either positive or negative) between individual metabolites in the natural accessions was observed for both genetic- and metabolic-based analysis (Supplementary files 3 and 4). Taken together, results of the meta-analysis indicate genetic co-regulation of the biosynthesis of specific isoprenoids.

Next, we conducted hierarchical clustering, in which the correlation matrix was used as a measure of the distance between metabolites in the natural accessions and the mapping population. This clearly showed relationships between metabolite levels (Figure 11), which might reflect coupling(s) in their biosynthetic pathways (Figure 12). Thus, chlorophylls and carotenoids were the most closely related compounds (Figure 11), while phytosterols, plastoquinone, and Prens formed a separate cluster, which was also attracting the Dol cluster. The Dol cluster was, however, much more distant from the three other metabolites. The most distant cluster was comprised of tocopherols and it did not seem to correlate significantly with any other metabolite. A small disagreement between the trees deduced for the natural accessions or for the EstC mapping population was found concerning the location of phytosterols and plastoquinone vs. Prens (Figure 10B and Figure 11).

**Figure 11.**
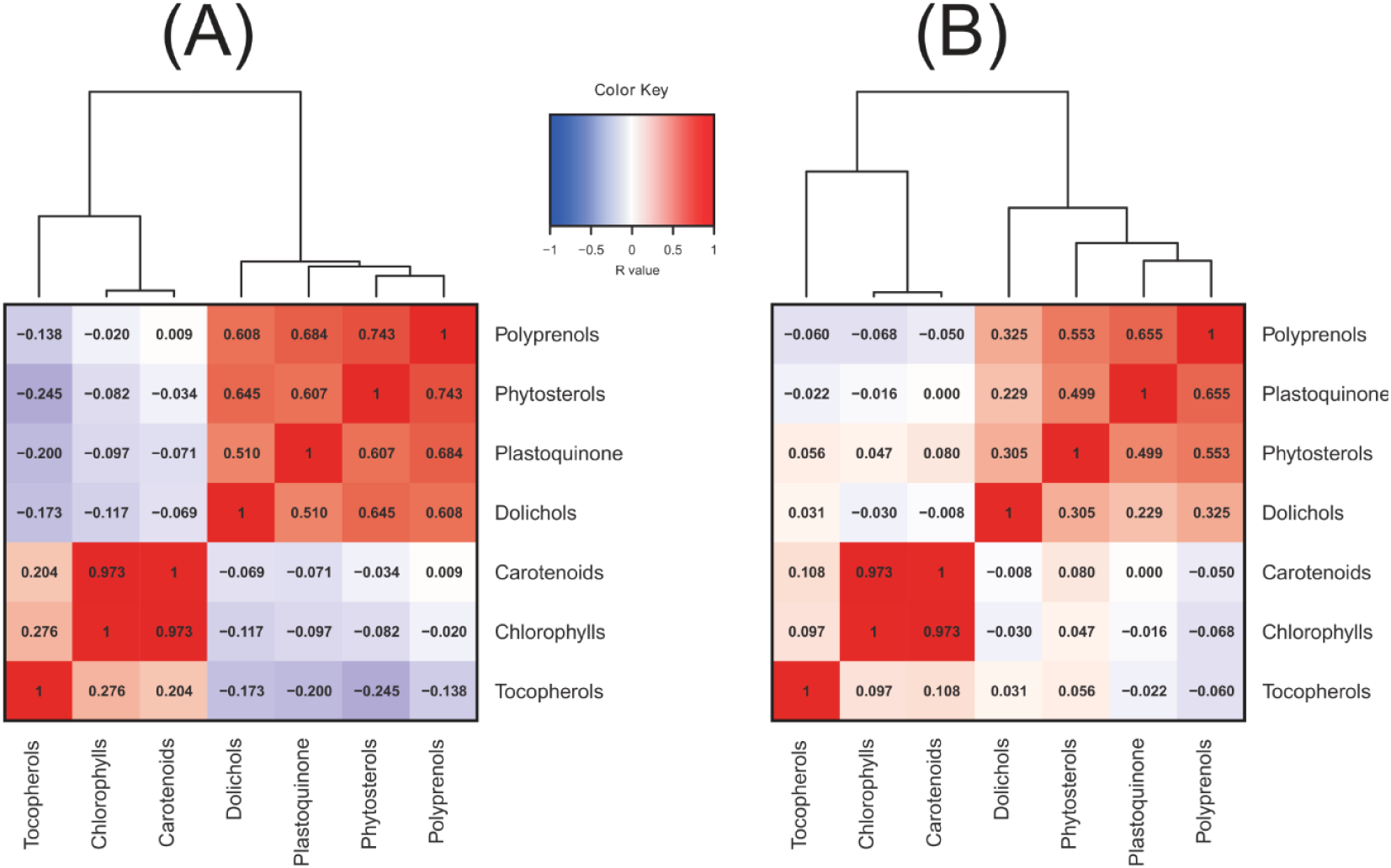
Dendrograms and corresponding heatmaps calculated for the accessions (A) and the mapping population (B). Hierarchical cluster analysis was performed for correlation matrixes (built of Spearman’s rank correlation coefficients, R) using the Lance–Williams dissimilarity update formula according to Ward’s clustering algorithm (Ward J, 1963) (see Materials and methods and Source Data 22).

**Figure 12.**
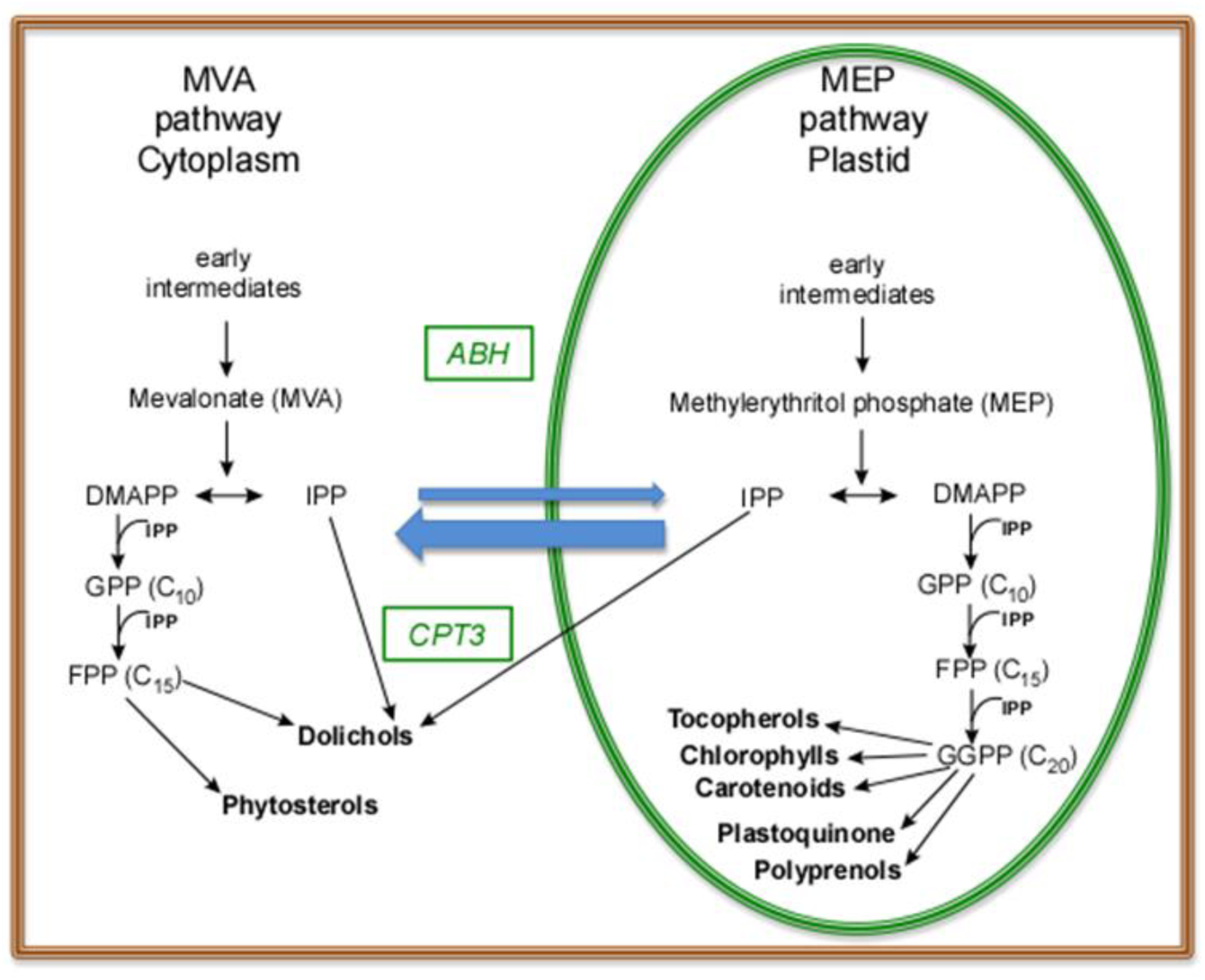
Biosynthetic routes leading to isoprenoids in a plant seedling cell; the involvement of the genes *cis*-prenyltransferase 3 (*CPT3*) and alpha-beta hydrolase (*ABH*) is indicated. Depicted are seven metabolites analyzed in this study. Two pathways, the mevalonate (MVA) and methylerythritol phosphate (MEP) pathways, are generating IPP in parallel, both contributing to particular isoprenoids (Hemmerlin et al., 2012; Akhtar et al., 2017; Jozwiak et al., 2017). Blue arrows illustrate the exchange of intermediates between the MVA and MEP pathways. Abbreviations: DMAPP-dimethyallyl diphosphate, FPP-farnesyl diphosphate, GPP-geranyl diphosphate, GGPP-geranylgeranyl diphosphate, IPP-isopentenyl diphosphate.

## DISCUSSION

### Genetic basis of Dol accumulation

A broad data set presented here shows that isoprenoid levels in Arabidopsis accessions are highly diverse. Estimated heritabilities from GWAS are non-zero for the analyzed traits, suggesting the existence of genetic factors underlying their accumulation. Our results also demonstrate the usefulness of the natural variation of Arabidopsis for identifying new loci underlying specific phenotypic traits.

It is intriguing that we could detect QTLs for four different compounds: Prens, Dols, chlorophylls, and carotenoids, while we found significant GWAS associations for three: phytosterols, plastoquinone, and Dols. Consequently, Dols are the only compounds where both approaches detected associations. Still, the reported QTL on chromosome 2 does not overlap with the GWAS results, which are located on chromosomes 1 and 3, respectively (summarized in Supplementary file 5). While, at a first glimpse, this lack of accordance might be disturbing, there could be many good reasons for it. It is well known that both methods have different power to detect associations (see Figure 4 in Weigel and Nordborg, 2015). For example, on chromosome 1, we identified a significant GWAS association for three different compounds, but we detected no corresponding QTL in the mapping population despite the fact that the associated polymorphism segregates in the AI-RIL population. The three traits for which this association is detected (the content of phytosterols, plastoquinone, and Dols) show a strong genetic correlation, so one would expect to find shared genetic factors that regulate all three traits, despite a slightly lesser phenotypic correlation of the traits. The associated sequence variant is located in the gene AT1G52450, which is thus an excellent candidate to modulate all three traits and would not have been found using QTL mapping alone. AT1G52450 is annotated to encode a ubiquitin carboxyl-terminal hydrolase (UCH)-related protein, while the neighboring gene AT1G52460 encodes an alpha-beta hydrolase, ABH (PubMed Gene database). Neither of these proteins has been characterized yet. Eukaryotic cells usually possess a family of UCHs (e.g., three in Arabidopsis) (Isono and Nagel, 2014) responsible for releasing ubiquitin (Ub) from ubiquitinated proteins. A tight balance between ubiquitination and deubiquitination is required for cellular survival since ubiquitin controls numerous bioactivities, such as protein degradation by the 26S proteasome, cell cycle regulation, signal transduction, or membrane trafficking. In turn, the ABH superfamily proteins are found across all domains of life. They are implicated in primary and secondary metabolism by serving highly diverse enzymatic activities, e.g., as esterases, thioesterases, lipases, proteases. Additionally, proteins with the α/β hydrolase fold function as receptors in the strigolactone, gibberellin and karrikin-smoke response pathways (Mindrebo et al., 2016 and references therein). In Arabidopsis, more than 600 proteins with ABH folds have been predicted by the InterPro database (Mitchell et al., 2019) with the majority remaining uncharacterized.

Taken together, hydrolytic enzymes, as ABH, encoded by AT1G52460, and/or UCH, encoded by AT1G52450, and putative ubiquitinating enzymes (respective genes detected by QTL, Supplementary file 1 and 2) might constitute a regulatory loop controlling isoprenoid biosynthesis in eukaryotic cells. Interestingly, both *ABH* and *UCH* show a high dN/dS ratio (ratio of nonsynonymous to synonymous divergence) in the Arabidopsis population, arguing for strong selection on these genes (see Supplementary file 6). Further studies are needed to identify the cellular target(s) of AT1G52460 and the mechanisms underlying its involvement in the metabolism of Dol, phytosterol, and plastoquinone.

It is worth noting that in previous reports, the AT1G52460 gene was identified as one of the maternally expressed imprinted genes (MEGs) that was shown to be predominantly expressed from maternal alleles in reciprocal crosses (Wolff et al., 2011). Notably, the AT1G52460 was among the MEGs (∼30% of all the MEGs tested in that study) for which authors reported a dN/dS value greater than one (Wolff et al., 2011). The dN/dS value can be used to measure the rate of molecular evolution of genes (Warren et al., 2010); therefore the results of Wolff et al. (2011) provide particularly strong evidence for the fast evolution of AT1G52460. Taking into account that we detected only heterozygotic lines for the AT1G52460 gene, we consider that a loss-of-function allele may lead to a lethal phenotype. This finding could be particularly important and it deserves further investigation since very few imprinted genes have been confirmed in plants and even fewer of them have been functionally investigated (He et al., 2017).

The confidence intervals of the detected QTLs include hundreds of different genes. This is within the typical mapping resolution of QTL studies but leads to the problem of prioritizing candidate genes. The most promising gene identified in the QTL analysis, AT2G17570 (*CPT3*), is a long searched enzyme responsible for backbone synthesis for the major family of dolichols in Arabidopsis, with Dol-15 and Dol-16 dominating. Interestingly, the different product specificity of the Arabidopsis enzymes CPT3, CPT6 (which produces *in planta* a single Dol-7 (Surmacz et al., 2014)) and the recently characterized CPT1 (producing a family of Dols with Dol-21 dominating) suggests that the particular AtCPTs play dedicated, non-redundant roles in isoprenoid synthesis in Arabidopsis tissues. For further comments regarding CPT3 see also Supplementary file 6.

Despite the fact that no overlapping associations have been found for the GWAS and QTL results, one can try, using the GWAS results, to prioritize candidate genes in the QTL interval. In the confidence interval of the detected QTL for Dol on chromosome 2, we could analyze 6,668 independent segregating polymorphisms with a minor allele frequency greater than 5%. None of these reached the genome-wide significance threshold; the most significant polymorphism had a p-value of 4.88^*^10^^^-6 and was located in the proximity of AT2G17570, which encodes CPT3. Although this score is marginal, it is locally significant, if we restrict our analysis to sequence variants within the QTL region. So the combined results of GWAS and QTL strongly indicate that *CPT3* is the gene underlying the detected QTL for Dol, despite the plethora of other tempting candidate genes. Detailed SNP analyses of *CPT3* revealed that this gene shows a high amount of variation with a total number of 30 non-synonymous substitutions and 5 alternative starts and 1 premature stop codon in the Arabidopsis population (Supplementary file 6).

### Trait correlations

Strong correlations between the levels of particular metabolites (Figure 10 and Figure 11) probably mirror common regulatory mechanisms responsible for their formation. Despite the fact that all studied metabolites are derived from the isoprenoid pathway, their clustering reflects the complexity of this pathway (Figure 12) and supports the notion of channeling of substrates and intermediates described earlier as ‘metabolons’ (Newman and Chappell, 1999). Thus, the tight (‘perfect’) association of carotenoids and chlorophylls is in agreement with their intimately linked function in photosynthesis, as well as their biosynthetic origin from a common isoprenoid precursor, geranylgeranyl diphosphate (GGPP), and the plastidial localization of their biosynthesis (Figure 12). Moreover, IPP molecules required for the synthesis of carotenoids, chlorophylls, and plastoquinone are thought to be derived mostly from the methylerythritol phosphate (MEP) pathway and several, but not all, steps of their synthesis are located in plastids too. Prens seem to cluster more closely with plastoquinone than with Dols (Figure 11), despite the high similarity of Pren and Dol structure. Interestingly, a growing body of information suggests that the biosynthetic routes for Prens and Dols in plants are different. Prens are synthesized in plastids (Akhtar et al., 2017), probably from MEP-derived IPP, while Dols are synthesized consecutively in plastids and in the cytoplasm (Skorupinska-Tudek et al., 2008; Jozwiak et al., 2017) with the concomitant involvement of the MEP and MVA pathways (Figure 12). Thus, the assignment of Prens and Dols to distinct clusters and the calculated dendrogram are in line with the described above differences in their biosynthetic routes.

## CONCLUSIONS

In this study several candidate genes for potential new factors that might regulate polyisoprenoid accumulation have been identified. The regulation of isoprenoid pathways is complex, but using the combination of both GWAS and QTL it is possible to prioritize the underlying genes.

Understanding of the mechanisms of Dol synthesis/accumulation in eukaryotes is important since the shortage of dolichol/dolichyl phosphate results in serious defects in all studied organisms, most probably caused by defective protein glycosylation. In plants, it is lethal due to male sterility (Jozwiak et al., 2015; Lindner et al., 2015) while in humans mutations in genes encoding enzymes involved in Dol/DolP synthesis lead to rare genetic disorders collectively called Congenital Disorders of Glycosylation (CDG type I); supplementation of the diet with plant tissues that can be utilized as a source of dolichol/dolichyl phosphate has been suggested (summarized in Buczkowska et al., 2015). The identification of genes regulating the synthesis/accumulation of Dols – such as the here detected *CPT3* and *ABH* – opens a perspective for the manipulation of Dol content in plants and consequently makes it feasible to think of constructing plants with increased Dol content.

## MATERIALS AND METHODS

### Plant materials

*Arabidopsis thaliana* accessions used in this study are listed in the Supporting Information (Supplementary file 7). All accessions were obtained from the stock center NASC (http://arabidopsis.info/).

A population of advanced intercross recombinant inbred lines (AI-RIL, EstC) was obtained after crossing of the Est-1 (Estland) and Col-0 (Columbia) accessions (Balasubramanian et al., 2009). All lines were kindly provided by Maarten Koornneef from Max Planck Institute for Plant Breeding Research in Cologne, Germany. The EstC mapping population with all sequence variant database is available at the NASC under the stock number CS39389.

For miRNA-mediated knockdown of the *CPT3* gene, two pairs of primers specific to amiRNA and amiRNA^*^ targeting the gene were designed using the Web MicroRNA Designer WMD3. The vector pRS300 was used as a template for subsequent PCR amplification and replacement of the endogenous miR319a and miR319a^*^ sequences with appropriate amiRNA and amiRNA^*^ of *CPT3* as described in the website protocol wmd3.weigelworld.org (Ossowski Stephan, Fitz Joffrey, Schwab Rebecca, Riester Markus and Weigel Detlef, personal communication). The obtained stem-loop was used as a template for PCR to generate the 454 bp fragment with a CACC overhang at the 5′ end, which was used for directional cloning into the pENTR/D-TOPO vector system (Invitrogen). The recombination reaction from pENTR/D-TOPO to the pGWB602 binary vector was carried out with the Gateway LR clonase II system (Invitrogen). All primers used in the construction of the *AtCPT3* silencing vector are listed in Supplementary file 8. The obtained plasmid was introduced into *Agrobacterium tumefaciens* strain GV3101, which was then used to transform Arabidopsis (Col-0) by the floral dip method (Weigel and Glazebrook, 2002). T1 seeds were germinated on soil and transgenic plants were selected by spraying with 0.1% BASTA in the greenhouse. Spraying was performed one week after germination and was repeated two times at two-day intervals. Additionally, the plants that survived were verified by PCR.

AtCPT3-over-expressing lines (*CPT3-OE*) were generated using a 35S::AtCPT3 construct introduced into the *A. tumefaciens* GV3101 strain. Transformation of Arabidopsis (Col-0) plants was performed by the floral dip method (Weigel and Glazebrook, 2002). Transformant selection was performed as described previously (Surowiecki et al., 2019).

The T-DNA insertion mutant lines for AT1G52460, SALK_066806 and GK_823G12, were obtained from the Nottingham Arabidopsis Stock Center, their progeny were genotyped, and heterozygous lines were isolated.

### Growth conditions

Plants were grown in a growth chamber in a long day (16 h light) photoperiod at 22 ^°^C/18 ^°^C at day/night. The seeds were surface-sterilized by treatment with an aqueous solution of 5% calcium hypochlorite for 8 min, subsequently rinsed four times with sterile water and planted on plates. Before location in the growth chamber, plates with seeds were kept for 4 days at 4 ^°^C in darkness for stratification. The Arabidopsis accessions, the AI-RIL mapping population and the T-DNA insertion mutant lines were grown on Petri dishes on solid ½ Murashige-Skoog medium with vitamins (1L of medium contained 0.5 µg nicotinic acid, 0.5 µg pyridoxine, 0.1 µg thiamine, 2 µg glycine) and 0.8% agar. For each genotype analyzed (accessions, mapping population, T-DNA mutants), plants were cultivated in at least three biological replicates.

### Isolation of isoprenoids

Unless indicated otherwise, entire 3-week-old seedlings were used for the isolation of all isoprenoid compounds. To elucidate the correlation between polyisoprenoid content vs. *CPT3* transcript level, the Arabidopsis seedlings, leaves, and flowers were used. For chromatographic analysis of isoprenoids, either internal (Prens, Dols, and phytosterols) or external (plastoquinone and tocopherol) standards were employed.

#### Prens, Dols, and phytosterols

analyses were performed as described earlier with modifications (Gawarecka and Swiezewska 2014). Briefly, 3 g of fresh seedlings, supplemented with internal standards of Pren-14 (15 μg) and cholestenol (10 μg), were homogenized in 20 ml of chloroform/methanol solution (1/1, v/v) and extracted for 24 h at 25 ^°^C, lipids were subjected to alkaline hydrolysis, purified on silica gel columns, dissolved in isopropanol (final concentration 6 mg per 1 ml) and stored at -20 ^°^C until used.

#### Plastoquinone

0.5 g of seedlings was used. Isolation procedure was as described above, but the hydrolysis step was omitted and the samples were protected from light.

#### Chlorophylls and carotenoids

0.2 g of seedlings was homogenized in acetone, extracted for 24 h at 25 ^°^C, centrifuged (2500 × *g*) and the supernatant was directly subjected to spectrophotometric analyses. All isolation steps were performed in darkness.

#### Tocopherol

3 g of seedlings were homogenized in 6 ml ethanol and extracted for 24 h at 25 ^°^C, the sample was supplemented with 4 ml of water and 3 ml of a mixture of hexane/dichloromethane (9/1, v/v) to separate the phases. Water phase was re-extracted 3 times with 3 ml of hexane/dichloromethane, organic phases were pooled and evaporated, lipids were dissolved in 8 ml of dichloromethane and analyzed directly. During the preparation, samples were protected from light.

### HPLC/UV analyses of polyisoprenoids and plastoquinone

HPLC/UV analyses of polyisoprenoids were performed as described earlier (Gawarecka and Swiezewska, 2014) with modifications. Briefly, a Waters dual λ absorbance detector and a 4.6 × 75 mm ZORBAX XBD-C18 (3.5 µm) column (Agilent, USA) were used. The applied solvent system was (A) methanol/water (9/1, v/v), (B) methanol/hexane/propan-2-ol (2/1/1, v/v/v) and a gradient program was from 100 – 35% A for 3 min, 35 – 0% A for 7 min, 100% B for 8 min. Qualitative analyses were performed using external standards – mixtures of Prens (Pren-9, -11, …, -23, -25) and Dols (Dol-16 to -21) – while quantitative analyses were performed using the internal standard, Prenol-14. All standards were from the Collection of Polyprenols, IBB PAS, Warsaw, Poland.

HPLC/UV analyses of plastoquinone were performed using the above protocol with a slightly modified gradient: 100 – 35% A for 3 min, 35 – 0% A for 7 min, 100% B for 5 min.

### GC/FID analysis of phytosterols and tocopherols

GC analysis was performed employing an Agilent Technologies, 7890A apparatus equipped with a split/splitless injector and an FID detector with an HP-5 column (J & W Scientific Columns, Agilent Technologies) 30 m × 0.32 mm and 0.25 µm film thickness.

Phytosterols were analyzed as described previously (Jozwiak et al., 2013). Signals were identified by comparison with external standards (Sigma-Aldrich-Fluka, Poznan). The following compounds were identified in plant samples: campesterol, stigmasterol, β-sitosterol, stigmast-4,22-dien-3one, stigmast-4en-3-one, brassicasterol, β-sitostanol, cholesterol. Total content of phytosterols was used for further analyses.

Tocopherols were analyzed as described previously (Kadioglu et al., 2009). Signals of tocopherol α, δ and γwere identified by comparison with external standards (a kind gift of Prof. Gustav Dallner, University of Stockholm). Total content of tocopherols was used for further analyses.

### Spectrophotometric analyses of chlorophylls and carotenoids

Chlorophylls and carotenoids were analyzed as described earlier (Lichtenthaler and Buschman, 2001). All analyses were performed in triplicate (three independent biological replicates). The amounts of all isoprenoid compounds were expressed as μg per g of fresh weight.

### Complementation of the yeast *rer2*Δ mutant

To express *CPT3* and *LEW1* in *Saccharomyces cerevisiae* mutant cells (*rer2*Δ mutant: *rer2*::kanMX4 *ade2-101 ura3-52 his3-200 lys2-801*), coding sequences of *CPT3* and *LEW1* (AT1G11755) were subcloned into the pESC-URA yeast dual expression vector (Agilent, Santa Clara, CA, USA) according to the manufacturer’s protocol. Transformant selection and growth, as well as analyses of polyisoprenoid profile and CPY glycosylation status, were performed as described previously (Surowiecki et al., 2019).

### Subcellular localization and BiFC assays

For subcellular localization analysis of *35S::CPT3, A. tumefaciens* cells carrying the vectors CPT3-GFP and cd3-954 (ER-CFP, used as an organelle marker) were introduced into the abaxial side of *Nicotiana benthamiana* leaves. A BiFC assay was performed based on split EYFP. EYFP was fused to the C-terminus of CPT3 and the N-terminus of Lew1, resulting in the expression of CPT3:EYFPC and Lew1:EYFPN. CPT3:EYFPC was co-infiltrated with Lew1:EYFPN into the abaxial side of *N. benthamiana* leaves. A positive fluorescence signal (EYFP) is indicative of the restoration of EYFP due to the heterodimerization of CPT3 with Lew1.

The transient expression of CPT3, ER-CFP, and CPT3/Lew1-YFP fusion proteins was observed under a Nikon C1 confocal system built on TE2000E with 408, 488 and 543 nm laser excitations for CFP (450/35 nm emission filter) and GFP (515/30 nm emission filter), respectively.

### Statistical analyses

#### Quantitative genetic analyses

Mean values of at least three replicates were calculated for each isoprenoid compound measured, for each AI-RIL and each natural accession. These values were used in QTL mapping and GWAS. The broad sense heritability (H^2^) for isoprenoid accumulation for the AI-RIL population was estimated according to the formula: H^2^ = V_G_ /(V_G_ + V_E_), where VG is the among-genotype variance component and VE is the residual (error) variance. For GWAS heritability, estimates have been extracted from the mixed model accordingly.

#### QTL analyses in the AI-RIL population

All obtained phenotypical data were used in QTL mapping that was performed using R software (R Core Team, 2012, https://www.R-project.org/) with R/qtl package (Arends et al., 2010, Broman et al., 2003; http://www.rqtl.org/). Stepwise qtl function was used to detect multiple-QTL models (Broman, 2008, http://www.rqtl.org/tutorials/new_multiqtl.pdf). This function requires single-QTL genome scan to locate QTLs with the highest LOD scores, then the initial model is tested using arguments for additional QTLs and interactions between QTLs search, model refinement and backward elimination of each QTL detected back to the null model. Obtained QTL models were refined with the refineqtl function; any possible interactions between QTLs were verified by the addint function.

#### GWAS

Genome-wide association mapping was performed on measurements for 115 – 119 different natural accessions per phenotype. The phenotypic data are available at the AraPheno database (Seren et al., 2016). The genotypic data were based on whole-genome sequencing data (The 1001 Genomes Consortium, 2016) and covered 4,314,718 SNPs for the 119 accessions. GWAS was performed with a mixed model correcting for population structure in a two-step procedure, where first all polymorphisms were analyzed with a fast approximation (emmaX, Kang et al., 2010) and afterwards the top 1000 polymorphisms were reanalyzed with the correct full model. Only polymorphisms with a minor allele count greater than 5 are reported. The kinship structure has been calculated under the assumption of the infinitesimal model using all sequence variants with a minor allele frequency of more than 5% in the whole population. The analysis was performed in R (R Core Team, 2016). The R scripts used are available at https://github.com/arthurkorte/GWAS. The genotype data used for GWAS are available at the 1001 Genomes Project (www.1001genomes.org).

#### Correlation analyzes of isoprenoid accumulation - a statistical meta-analysis

All correlation analyses were performed with the aid of R version 3.3.0 (R Core Team, 2016, https://www.R-project.org/) using the outliers (Komsta, 2011, R package version 0.14, https://CRAN.R-project.org/package=outliers) and the gplots (Warnes et al., 2016, R package version 3.0.1, https://CRAN.R-project.org/package=gplots). The significance level α of 0.001 was assumed in all statistical tests.

Although for each accession the level of each metabolite was measured in triplicate, the values thus obtained were analyzed separately, as indicated by the number of experimental points in the respective figures (which equals three times the number of accessions). Means were not calculated, and this approach was employed deliberately to avoid the problem of adjusting and weighing mean values and to allow testing for outliers among single replicates instead of among mean values.

The Shapiro-Wilk test (Shapiro and Wilk 1965) was used to assess the agreement of isoprenoid content in the populations with the Gaussian distribution. Since, even after filtering out of extreme values with the Grubbs’ test for outliers (Grubbs 1950), a vast majority of the distributions were found non-Gaussian, further analyses were performed using non-parametric methods. Consequently, a correlation matrix for the seven investigated isoprenoids was calculated accordingly to the Spearman’s rank correlation coefficients (Spearman 1904).

A hierarchical cluster analysis of the correlation matrix was performed according to the Ward criterion (Ward 1963).

### Selection of candidate genes from chosen QTL intervals

We selected one QTL for Dol (DOL1) and three QTLs associated with Pren accumulation (PRE1, PRE2, PRE3) for further *in silico* analyses. The selected intervals were characterized by the highest percentage of phenotypic variance explained by each QTL and the highest LOD (logarithm of the odds) score values linked with the lowest number of loci (Table 2). The positional candidate genes within QTL confidence intervals were extracted from the Araport11 Annotation (www.araport.org). Firstly, we checked the annotated functions for all genes located in the selected QTL intervals by analyzing available databases and literature data for the isoprenoid biosynthetic pathways (TAIR, http://www.arabidopsis.org; PubMed, https://www.ncbi.nlm.nih.gov/pubmed). In this way, we obtained lists of candidate genes putatively considered to be involved in the regulation of Pren and Dol biosynthesis/accumulation (Supplementary file 1 and 2). Subsequently, we performed *in silico* analyses focused on tissue distribution and expression levels of the selected genes using data from the Arabidopsis eFP Browser 2.0 database (http://bar.utoronto.ca). This procedure allowed us to generate four sets of genes – three for Pren (Supplementary file 1) and one for Dol (Supplementary file 2). Detailed SNP analyses of At2G17570 (*CPT3*), AT1G52450 (*UCHs*), and AT1G52460 (*ABH*) sequences (Supplementary file 6) in the Arabidopsis population were extracted from the Arabidopsis 1001 genomes data using a custom R script.

### Quantitative real-time PCR analysis

Total RNA from Col-0, Stw-0, and Or-0 seedlings (1-, 2-, and 3-week-old) and leaves (4-, 5-, and 6-week-old plants) was isolated and purified using RNeasy Plant Mini Kit (Qiagen) following the manufacturer’s instructions. RNA concentration and purity were verified using a NanoDrop™ 1000 Spectrophotometr (Thermo Scientific, Walthman, MA). RNA was treated with RNase-free DNase I (Thermo Scientific) according to the manufacturer’s instructions. 160 ng RNA per each sample was used for first-strand synthesis using SuperScript™ II First-Strand Synthesis System for RT-PCR (Thermo Scientific) and oligo-dT primers according to the manufacturer’s procedure. 2 µl of cDNA was used for real-time PCR analysis of *AtCPT3* expression, using 0.6 µl each of gene-specific primers (5′-GCGCTTATGTCGATGCTG-3′-F; 5′-CAGACTCAACCTCCTCAGG-3′-R) in a total volume of 20 µl of Luminaris HiGreen High ROX qPCR Master Mix (Thermo Scientific) in a real-time thermal cycler STEPOnePlus (A&B Applied Biosystems, Waltham, MA) as instructed.

## ACKNOWLEDGEMENTS

We would like to express our gratitude to Professor Maarten Koornneef for providing the AI-RILs seeds used in this study. We also would like to thank Dr Agata Lipko for initial characterization of mutant lines and Rafał Banasiuk for help with preparing high-quality figures. Dr Marta Hoffman-Sommer is kindly acknowledged for help with preparation of the manuscript. This research was supported by grants from the National Science Centre of Poland [UMO-2014/15/N/NZ3/04316] (KG), [UMO-2018/29/B/NZ3/01033] (ES), and [UMO-2014/15/B/NZ2/01073] (AI), and the National Research Foundation (NRF) of Korea [NRF-2017R1A2B3009624] (JHA).

## Figure Supplements

**Figure 2-figure supplement 1**. GC/FID chromatogram of phytosterols from Arabidopsis Col-0 seedlings.

**Figure 2-figure supplement 2**. HPLC/UV chromatogram of plastoquinone from Arabidopsis Col-0 seedlings.

**Figure 2-figure supplement 3**. GC/FID chromatogram of tocopherols from Arabidopsis Col-0 seedlings.

**Figure 3-figure supplement 1**. Content of selected isoprenoids in the seedlings of Arabidopsis accessions.

**Figure 4-figure supplement 1**. Frequency distribution of the content of (A) chlorophylls, (B) carotenoids, (C) phytosterols, (D) plastoquinone and (E) tocopherols in the seedlings of AI-RILs and their parental lines, Col-0 and Est-1.

**Figure 8-figure supplement 1**. Manhattan plot of genome-wide association results for polyprenols, chlorophylls and tocopherols.

**Figure 9-figure supplement 1**. Phenotypic appearance of 4-week-old detached leaves of AT1G52460-deficient (SALK_066806, *abh* heterozygous mutant) and wild-type plants (Col-0) grown in soil.

**Figure 10-figure supplement 1**. Cumulative distributions (CDF) of the content of seven studied metabolites analyzed in the seedlings of Arabidopsis accessions and AI-RILs.

## Supplementary Files

**Supplementary file 1**. Candidate genes potentially involved in polyprenol accumulation, selected from the mapped intervals (a) PRE1, (b) PRE2 and (c) PRE3.

**Supplementary file 2**. Candidate genes potentially involved in dolichol accumulation, selected from the mapped QTL interval DOL1.

**Supplementary file 3**. Genetic correlations between metabolite levels.

**Supplementary file 4**. Metabolic data-based correlations between metabolite levels.

**Supplementary file 5**. Summary of candidate genes involved in accumulation of Dol, plastoquinone, phytosterols and Pren – comparison of QTL and GWAS approaches.

**Supplementary file 6**. Detailed SNP analysis of the At2G17570 (*CPT3*), AT1G52450 (*UCH*) and AT1G52460 (*alpha-beta hydrolase*) sequences in the Arabidopsis population.

**Supplementary file 7**. *Arabidopsis thaliana* accessions used in this study.

**Supplementary file 8**. Sequences of oligonucleotides used for the construction of the *AtCPT3* silencing vector.

## Source Data

Figure 2–**Source Data 1** (xlsx file format)

Figure 2–figure supplement 2–**Source Data 2** (xlsx file format)

Figure 2–figure supplement 3–**Source Data 3** (xlsm file format)

Figure 3, Figure 3–figure supplement 1–**Source Data 4** (xlsx file format)

Figure 4, Figure 4–figure supplement 1, Table 1–**Source Data 5** (xlsx file format)

Figure 5A, Table 2–**Source Data 6** (csv file format)

Figure 5B, Table 2–**Source Data 7** (csv file format)

Figure 5C, Table 2–**Source Data 8** (csv file format)

Figure 5D, Table 2–**Source Data 9** (csv file format)

Figure 6AB–**Source Data 10** (xlsx file format)

Figure 6C–**Source Data 11** (xlsx file format)

Figure 6D_BLUE–**Source Data 12** (tif file format)

Figure 6D_GREEN–**Source Data 13** (tif file format)

Figure 6D_MERGE–**Source Data 14** (tif file format)

Figure 6E_BLUE–**Source Data 15** (tif file format)

Figure 6E_GREEN–**Source Data 16** (tif file format)

Figure 6E_MERGE–**Source Data 17** (tif file format)

Figure 7A–**Source Data 18** (xlsx file format)

Figure 7B–**Source Data 19** (jpg file format)

Figure 8, Figure 8–figure supplement 1–**Source Data 20** (csv file format)

Figure 9B–**Source Data 21** (xlsx file format)

Figure 10 and 11, Figure 10-figure supplement 1–**Source Data 22** (txt file format)

**Figure 2-figure supplement 1.**
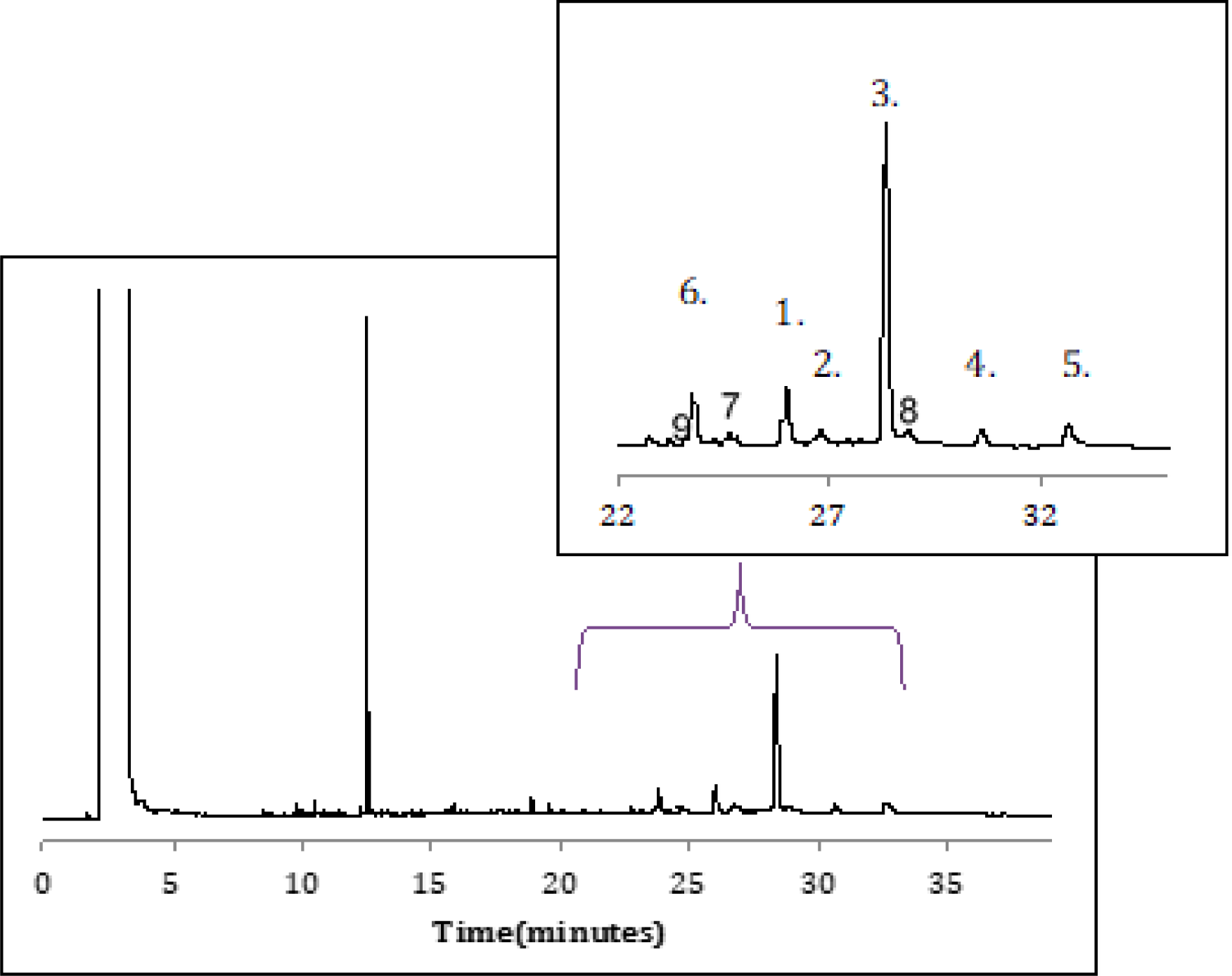
GC/FID chromatogram of phytosterols from Arabidopsis Col-0 seedlings. 1. campesterol; 2. stigmasterol; 3. β-sitosterol; 4. stigmast-4,22-dien-3one; 5. stigmast-4en-3-one; 6. cholestanol – internal standard; 7. brassicasterol; 8. β-sitostanol 9. cholesterol. The same profile of phytosterols was recorded for all analyzed accessions. Inset presents the magnified region of chromatogram. Indicated signals (1-5 and 7-9) were integrated to calculate the amount of phytosterols.

**Figure 2-figure supplement 2.**
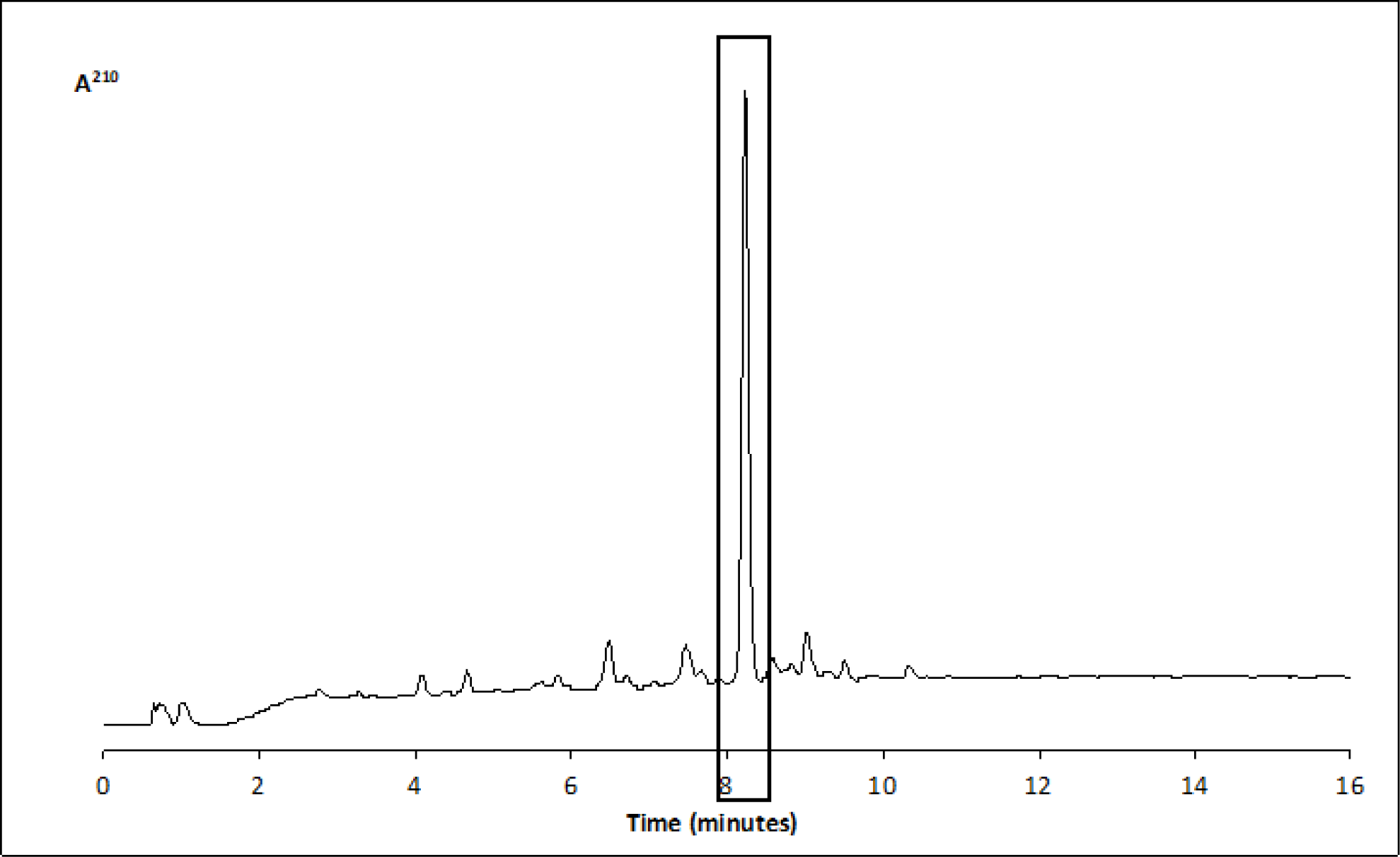
HPLC/UV chromatogram of plastoquinone of Arabidopsis Col-0 seedlings. The same lipid profile of was recorded for all analyzed accessions. Indicated signal was integrated to calculate the amount of PQ (Source Data 2).

**Figure 2-figure supplement 3.**
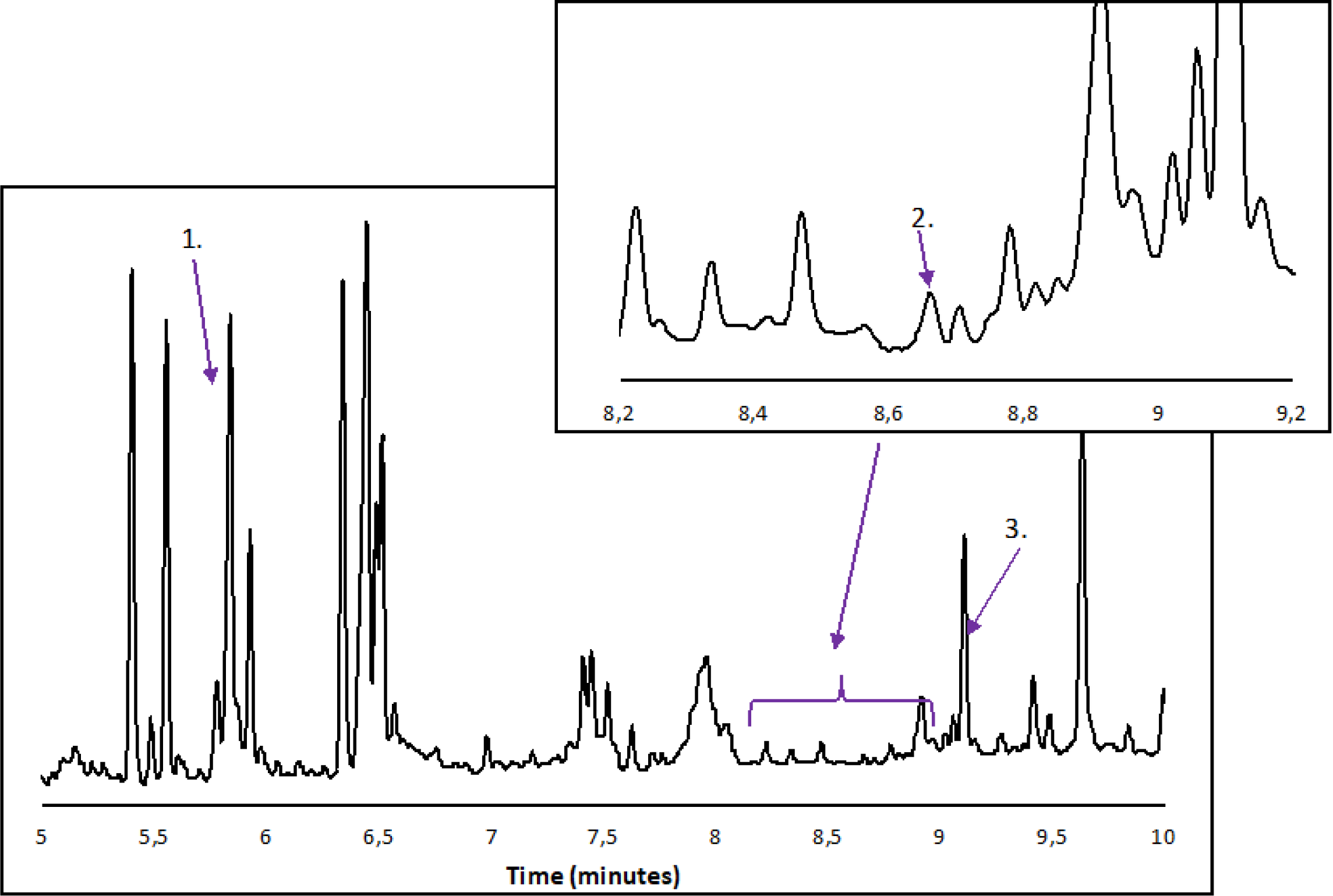
GC/FID chromatogram of tocopherols from Arabidopsis Col-0 seedlings. 1. γ - tocopherol; 2. δ - tocopherol (inset); 3. α - tocopherol of Arabidopsis Col-0 seedlings. The same lipid profile was observed for all analyzed accessions. Indicated signals (1-3) were integrated to calculate the amount of lipids (Source Data 3).

**Figure 3-figure supplement 1.**
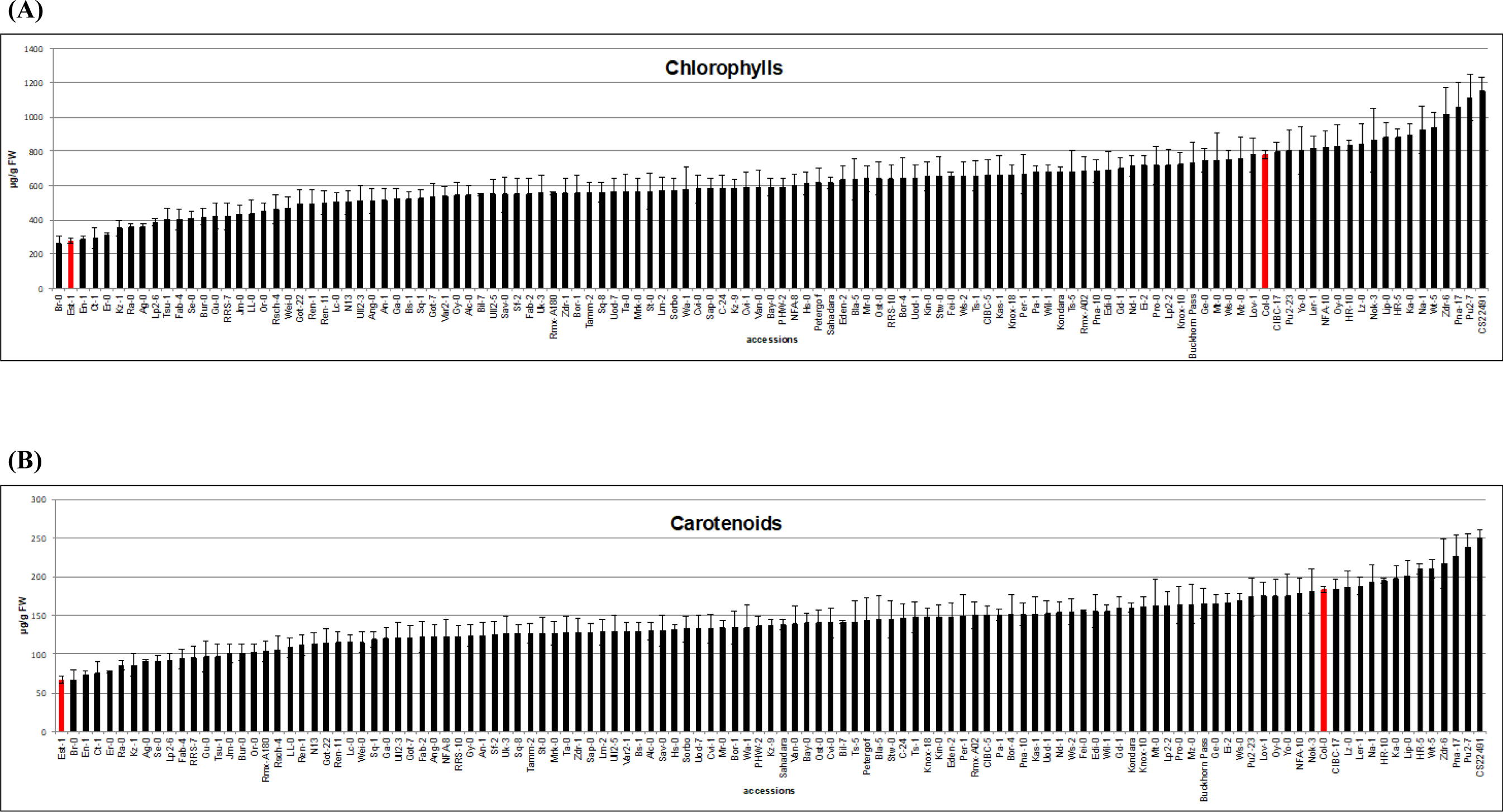
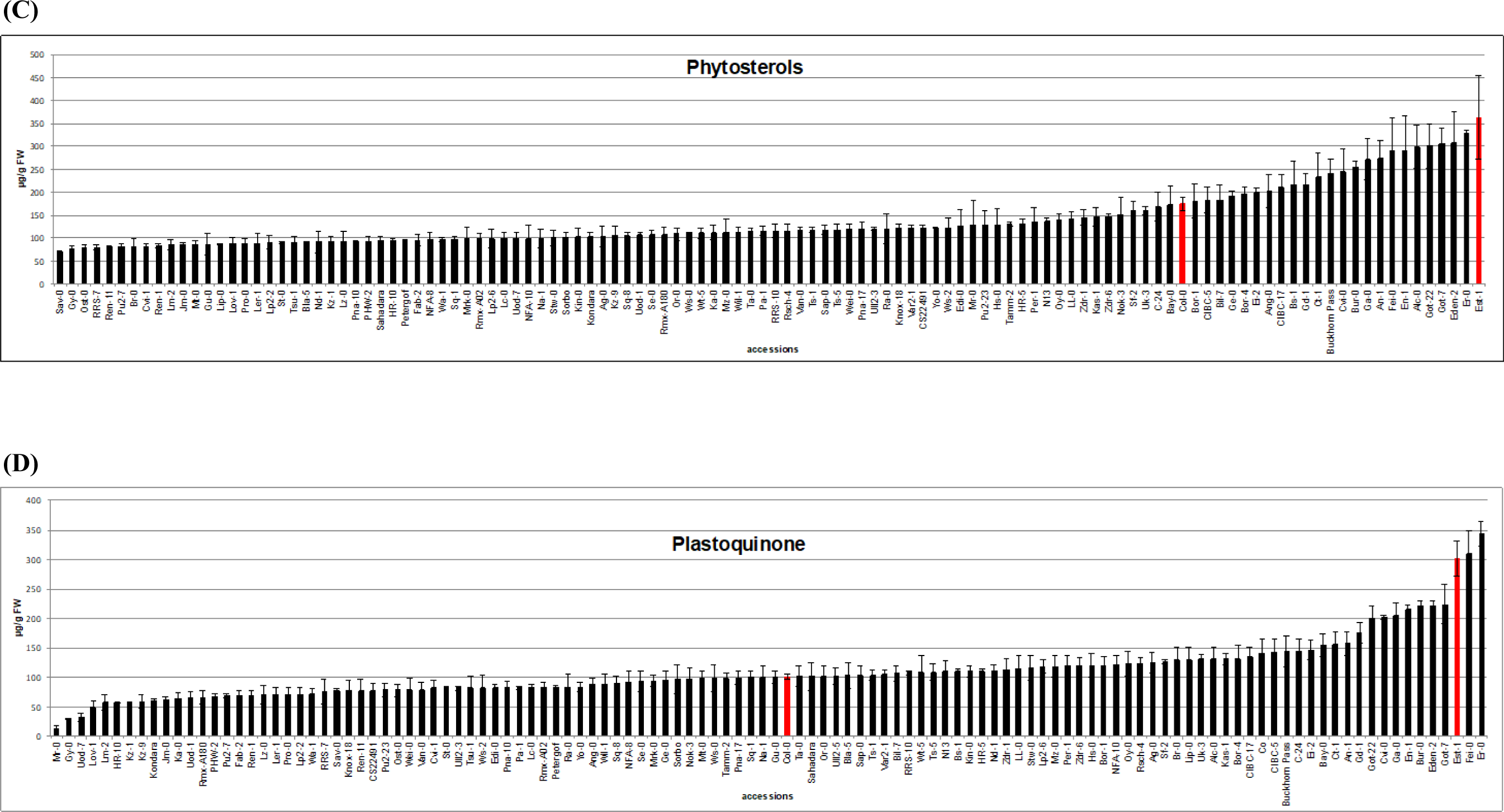
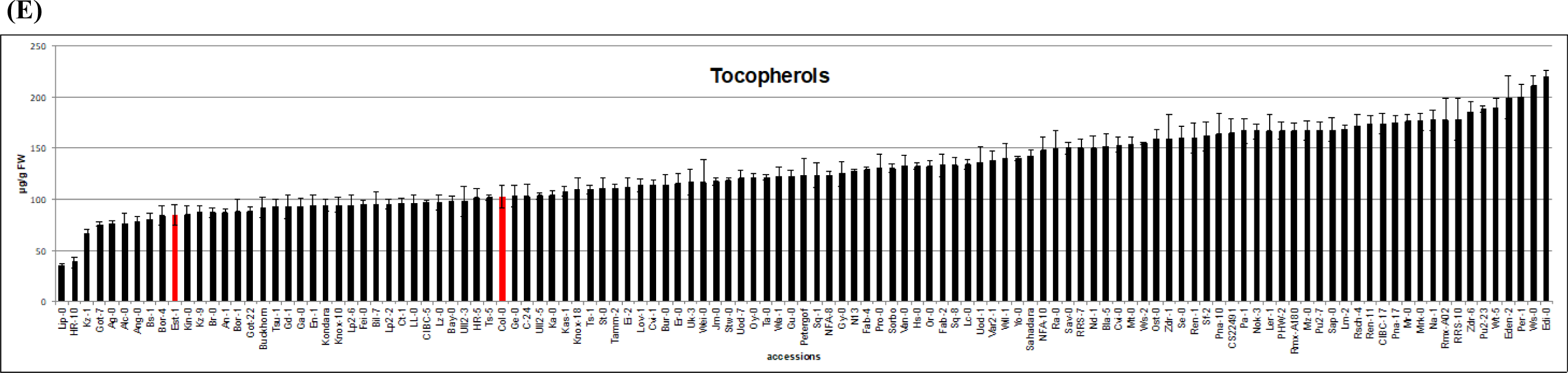
Content of selected isoprenoids in the seedlings of Arabidopsis accessions: (A) chlorophylls, (B) carotenoids, (C) phytosterols, (D) plastoquinone and (E) tocopherols. Bars presenting the content of particular isoprenoids in Col-0 and Est-1 are marked in red. All experiments were performed in triplicate, shown is mean ± SD. See Source Data 4.

**Figure 4-figure supplement 1.**
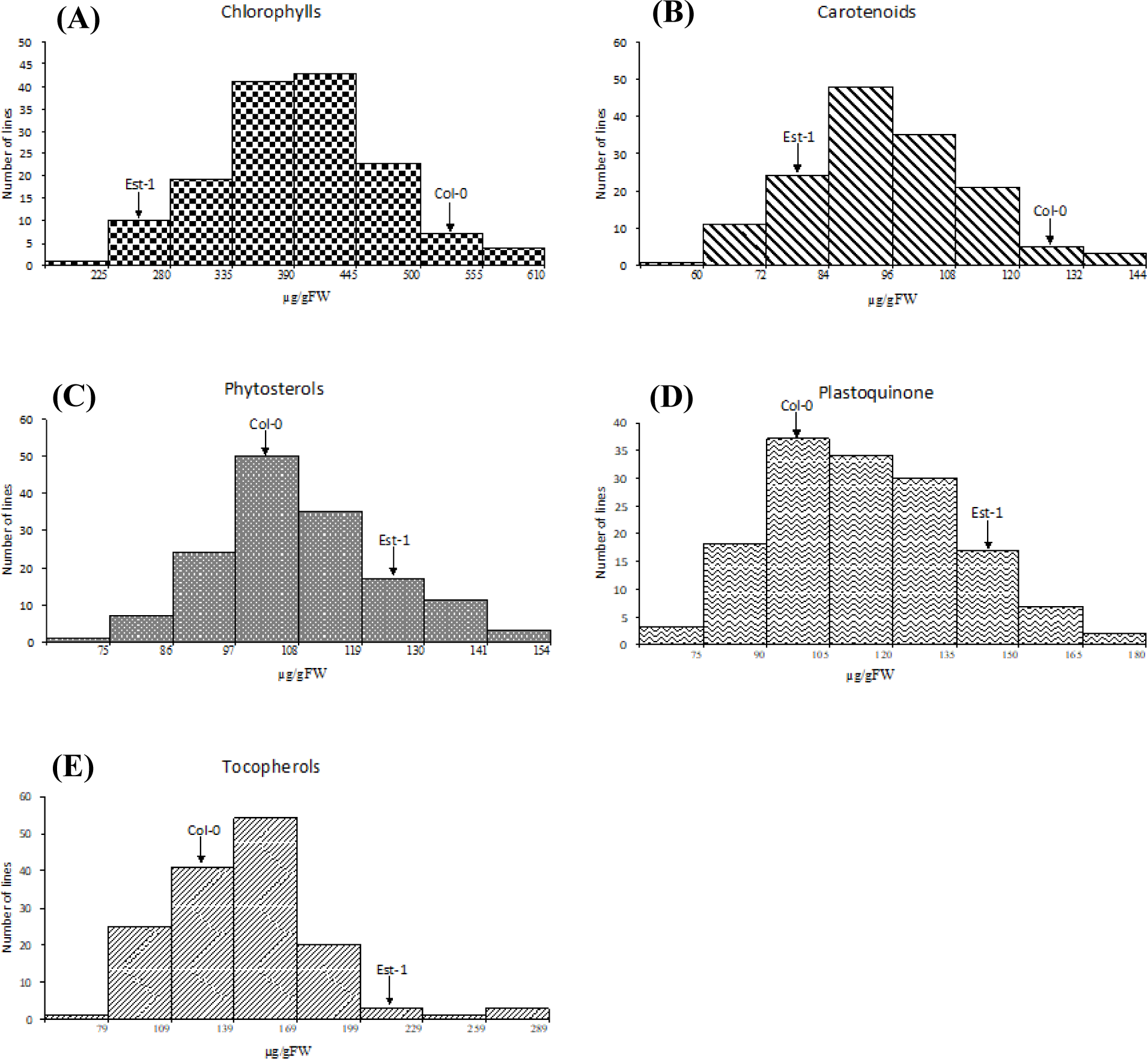
Frequency distribution of the content of (A) chlorophylls, (B) carotenoids, (C) phytosterols, (D) plastoquinone and (E) tocopherols in the seedlings of AI-RILs and their parental lines, Col-0 and Est-1. Each bar covers the indicated range of the content of a particular isoprenoid compound. See Source Data 5.

**Figure 8-figure supplement 1.**
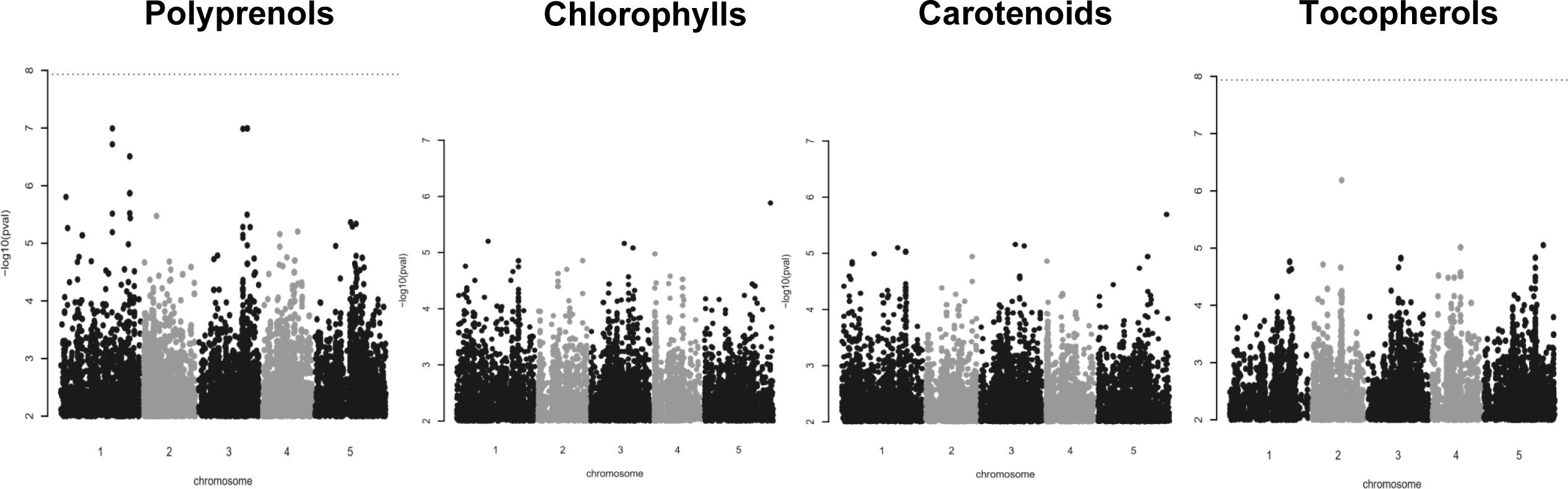
Manhattan plot of genome-wide association results for polyprenols, chlorophylls and tocopherols. The dotted horizontal lines indicate a significance level of 0.05 after Bonferroni correction for multiple testing. See Material and methods and Source Data 20.

**Figure 9-figure supplement 1.**
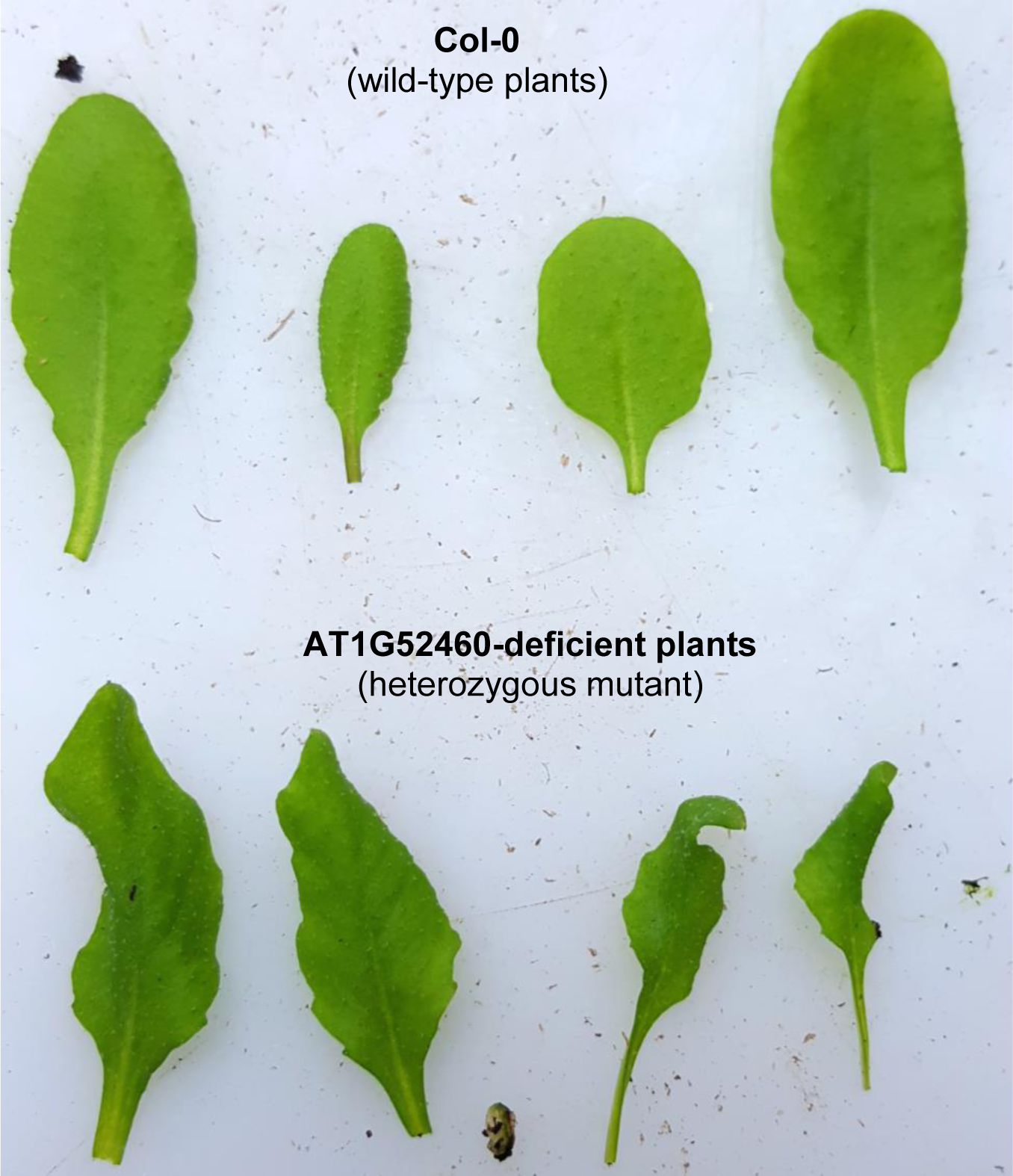
The phenotypic appearance of 4-week-old detached leaves of AT1G52460-deficient line (SALK_066806, *alpha-beta hydrolase*, heterozygous mutant) and wild-type (Col-0) plants grown in soil.

**Figure 10-figure supplement 1.**
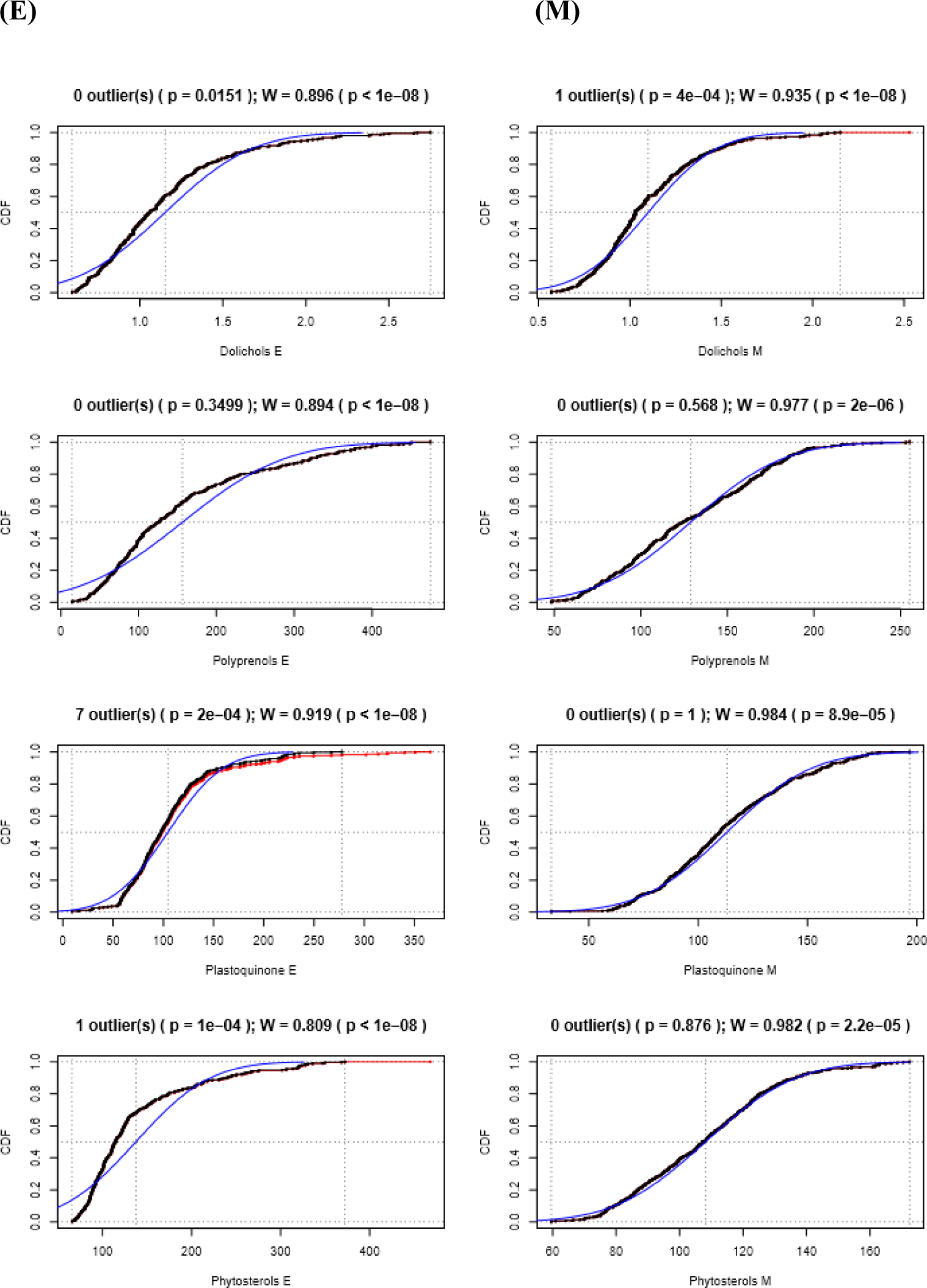
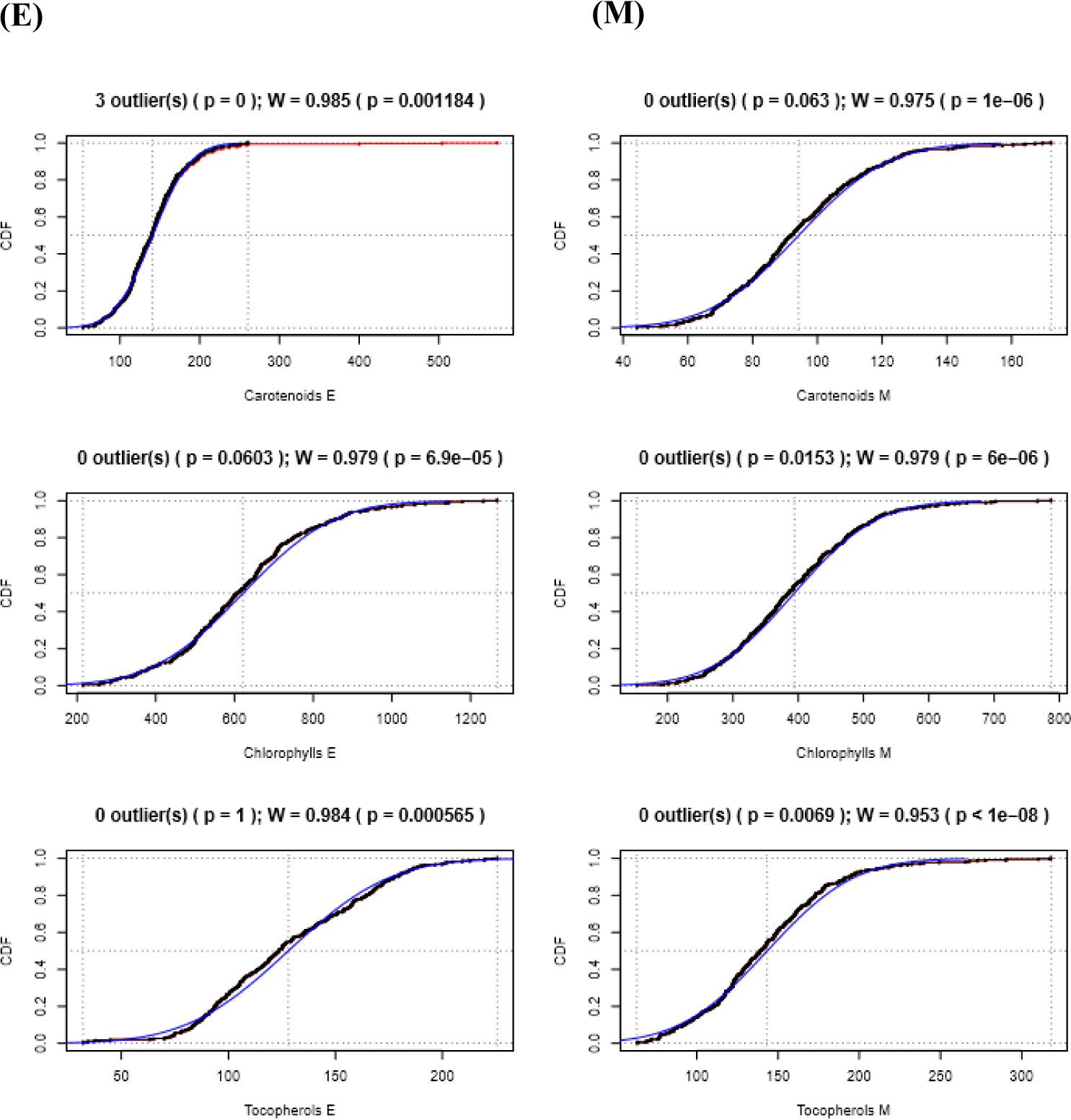
Cumulative distributions (CDF) of the content of seven studied metabolites analyzed in the seedlings of Arabidopsis accessions (E) and AI-RILs (M) (left and right column, respectively). Each set of data, presented in a single panel, was analyzed to check for the presence of outliers (Grubbs test at significance level α=0.001), and for normal distribution of the data filtered out of outliers (Shapiro-Wilk test). Red markers follow original distributions, while black ones show the same data with outliers removed. Blue lines represent the CDF expected for the normal distribution. Short statistics for Grubbs (G, p) and Shapiro-Wilk (W, p) tests are shown above each panel. See Material and methods and Source Data 22.

